# Specimen-Dependent Sampling and Signal Limitations Govern the Effectiveness of Low-Magnification Super-Resolution in Cryo-EM

**DOI:** 10.64898/2026.09.08.750297

**Authors:** Chun-Hsiung Wang, Kuen-Phon Wu, Yuan-Chih Chang

**Affiliations:** Institute of Biological Chemistry, Academia Sinica, Taipei, Taiwan; Academia Sinica Cryo-EM Center, Academia Sinica, Taipei 11529, Taiwan; Institue of Biochemical Sciences, National Taiwan University, Taipei, 10617, Taiwan

## Abstract

Single-particle cryo-EM routinely delivers near-atomic structures; however, optimizing data collection requires a delicate balance among magnification, sampling bandwidth, and particle throughput. Although low-magnification super-resolution imaging can recover information beyond the physical Nyquist limit, its benefit varies substantially among specimens. This variability suggests that the effectiveness of super-resolution may depend on whether reconstruction is limited by detector sampling bandwidth or by the recoverable particle signal. However, the conditions that distinguish these two regimes remain poorly defined. Here, we systematically compared super-resolution and physical-pixel workflows at two magnifications using apoferritin (APO) and malate synthase G (MSG) as representative specimens with contrasting molecular size, symmetry, and image contrast. At low magnification, APO exhibited sampling-limited behavior, with super-resolution processing achieving 1.72 Å compared with 2.74 Å for physical-pixel processing. In contrast, MSG exhibited predominantly signal-limited behavior under the same conditions, yielding comparable resolutions of 3.05 Å and 2.98 Å for super-resolution and physical-pixel processing, respectively, despite the increased sampling bandwidth. These contrasting responses were further supported by particle-number saturation, per-particle motion correction, and Q-score analyses, which provided complementary evidence for the underlying sampling-limited and signal-limited regimes. Together, these results provide a practical framework for assessing whether reconstruction quality is predominantly constrained by sampling bandwidth or recoverable particle signal and offer a rational basis for balancing achievable resolution and particle throughput when selecting acquisition strategies.

## 1. Introduction

Single-particle cryo-electron microscopy (cryo-EM) now routinely enables near-atomic-resolution structure determination of macromolecular complexes. This progress has been driven by advances in direct electron detectors and computational methods that enable efficient frame acquisition, beam-induced motion correction, and increasingly effective reconstruction of structural information from noisy particle images (Li *et al*., 2013; McMullan *et al*., 2014; Scheres, 2012; Punjani *et al*., 2017). Achieving high resolution, however, still requires balancing several competing constraints during data collection: pixel sampling must be fine enough to capture high-frequency structural information, while field of view, beam exposure, and total collection time impose practical limits on data acquisition. Magnification sits at the center of this trade-off, simultaneously determining pixel sampling and field of view and thereby influencing the number of particles recorded per micrograph and overall data throughput. As a result, the choice of magnification is one of the most consequential decisions in cryo-EM experimental design.

One strategy for easing this trade-off is to collect data at lower magnification. Lower magnification expands the field of view, increasing the number of particles captured per micrograph and thereby improving collection throughput. Under conventional physical-pixel processing, however, this comes at the cost of coarser pixel sampling, which caps the maximum representable spatial frequency at the detector’s physical Nyquist limit. Super-resolution acquisition provides a potential means of mitigating this constraint by exploiting sub-pixel positional information recorded by modern direct electron detectors. Super-resolution processing can recover structural information beyond the physical Nyquist limit, in principle allowing low-magnification datasets to achieve resolutions otherwise accessible only at higher magnification (Li *et al*., 2013; Feathers *et al*., 2021; Sheng *et al*., 2022).

This benefit, however, has been demonstrated primarily using large, highly symmetric benchmark specimens (Feathers *et al*., 2021; Sheng *et al*., 2022). Such specimens provide favorable conditions for particle alignment and high-resolution reconstruction even when sampling density is reduced. Whether the same advantage extends to specimens with lower molecular mass, lower image contrast, or less favorable alignment properties remains unclear. More generally, the factors determining when low-magnification super-resolution improves reconstruction quality have not been systematically defined.

We hypothesized that the effectiveness of super-resolution processing is governed not simply by specimen size, but by the dominant information limitation governing reconstruction—whether resolution is capped by detector sampling bandwidth or by the recoverable signal within the particle images themselves. Under this view, super-resolution should provide a substantial benefit only when recoverable structural information extends beyond what physical-pixel sampling can represent. Conversely, when recoverable high-resolution information is limited by per-particle signal, image contrast, or alignment accuracy, expanding the sampling bandwidth should provide little additional gain (Wu & Lander, 2020; Herzik *et al*., 2017; Wentinck *et al*., 2022). Thus, the benefit of super-resolution is expected to depend on whether a reconstruction is predominantly sampling-limited or signal-limited under a given experimental condition.

To test this hypothesis, we compared super-resolution (Bin1) and physical-pixel (Bin2) processing workflows at two nominal magnifications, 130,000× and 64,000×, using apoferritin (APO, ∼500 kDa) and malate synthase G (MSG, ∼80 kDa) as specimens with contrasting size, symmetry, and image contrast. Within each dataset, Bin1 and Bin2 workflows were performed in parallel from the same movie stacks, allowing the effect of pixel sampling to be evaluated independently of differences in data collection. Reconstruction quality was assessed using global and local resolution, density-map features, and Q-score analysis, while particle-number saturation and per-particle motion correction analyses provided complementary evidence for the factors limiting reconstruction. This study provides a framework for assessing whether reconstruction quality is predominantly constrained by sampling bandwidth or recoverable particle signal under a given experimental condition, thereby providing a basis for specimen-aware acquisition and processing strategies.

## 2. Materials and Methods

### 2.1. Experimental design overview

The experimental design enabled direct comparison of super-resolution (Bin1) and physical-pixel (Bin2) reconstruction under matched data-acquisition conditions. Two specimens with distinct molecular and imaging properties were analyzed at nominal magnifications of 130,000× and 64,000×. For each dataset, Bin1 and Bin2 reconstructions were generated from the same movie stacks, allowing the effects of pixel sampling to be evaluated independently of data-collection variability. Reconstruction quality was evaluated using global and local resolution and density-map features, together with particle-number saturation, per-particle motion correction, and Q-score analyses.

### 2.2. Specimen preparation and vitrification

Mouse heavy-chain apoferritin (APO, ∼500 kDa) and *Escherichia coli* malate synthase G (MSG, ∼80 kDa) were used as representative specimens. Cryo-EM grids were rendered hydrophilic by glow discharge using a PELCO easiGlow system (Ted Pella) at a constant current of 20 mA. Grid vitrification was performed using a Vitrobot Mark IV (Thermo Fisher Scientific) at 4 °C and 100% relative humidity. A 4 μL aliquot of purified protein was applied to the grid, followed by blotting with filter paper and immediate plunge-freezing into liquid ethane cooled by liquid nitrogen. Vitrified grids were stored in liquid nitrogen until data collection. Detailed grid preparation and freezing parameters are summarized in Table S1.

### 2.3. Cryo-EM data acquisition

All cryo-EM data were collected on a 300-kV Titan Krios G3 transmission electron microscope (Thermo Fisher Scientific) equipped with an X-FEG electron source, a GIF BioQuantum energy filter operated with a 15-eV slit, and a K3 direct electron detector (Gatan). The detector was operated in super-resolution counting mode, and automated data acquisition was performed using EPU software (Thermo Fisher Scientific). For each specimen, datasets were acquired at nominal magnifications of 130,000× and 64,000×, corresponding to physical pixel sizes of 0.648 Å and 1.336 Å, respectively, and super-resolution pixel sizes of 0.324 Å and 0.668 Å. Movie stacks were recorded as unnormalized TIFF files. Detailed acquisition parameters are summarized in Table 1.

**Table 1.**
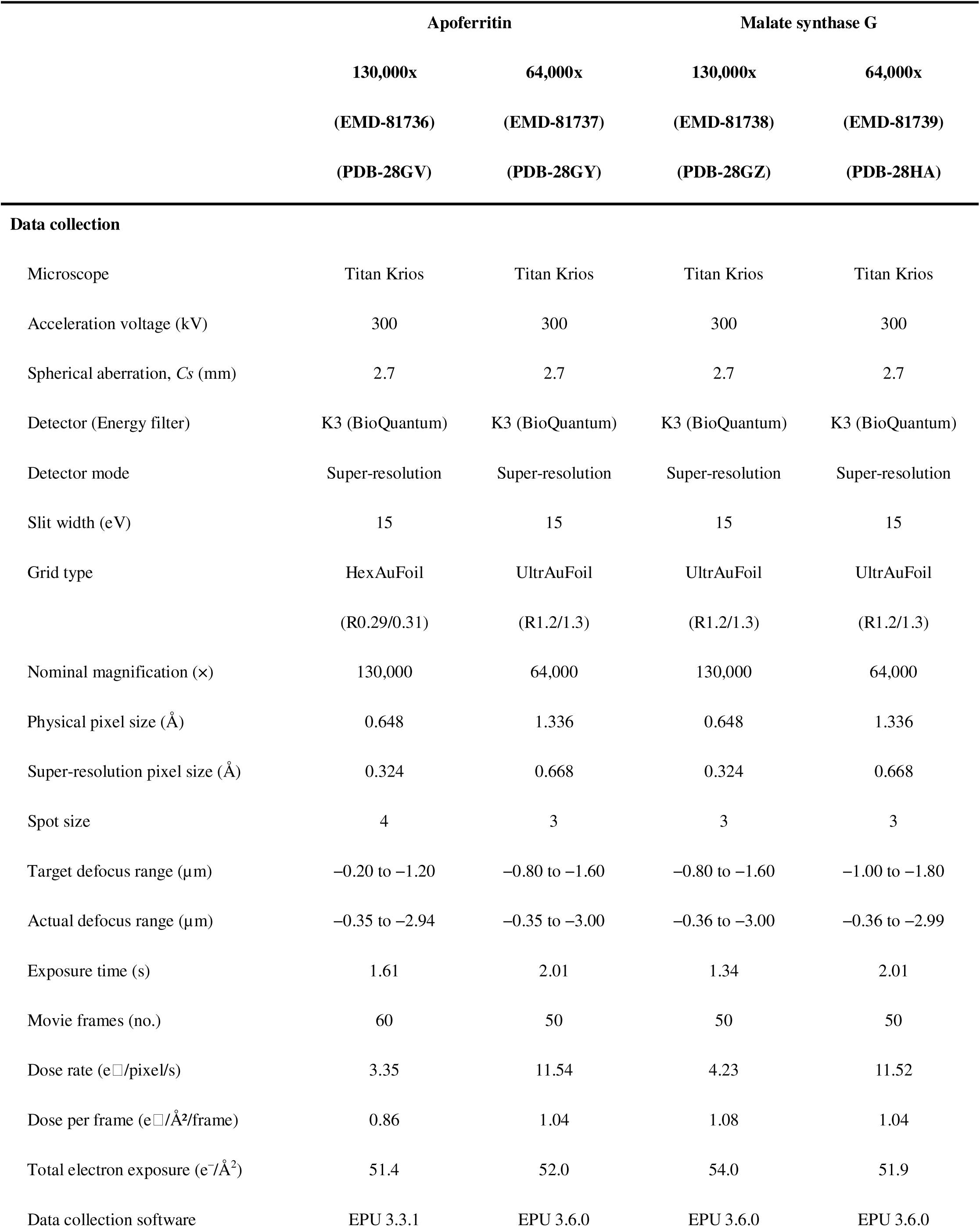
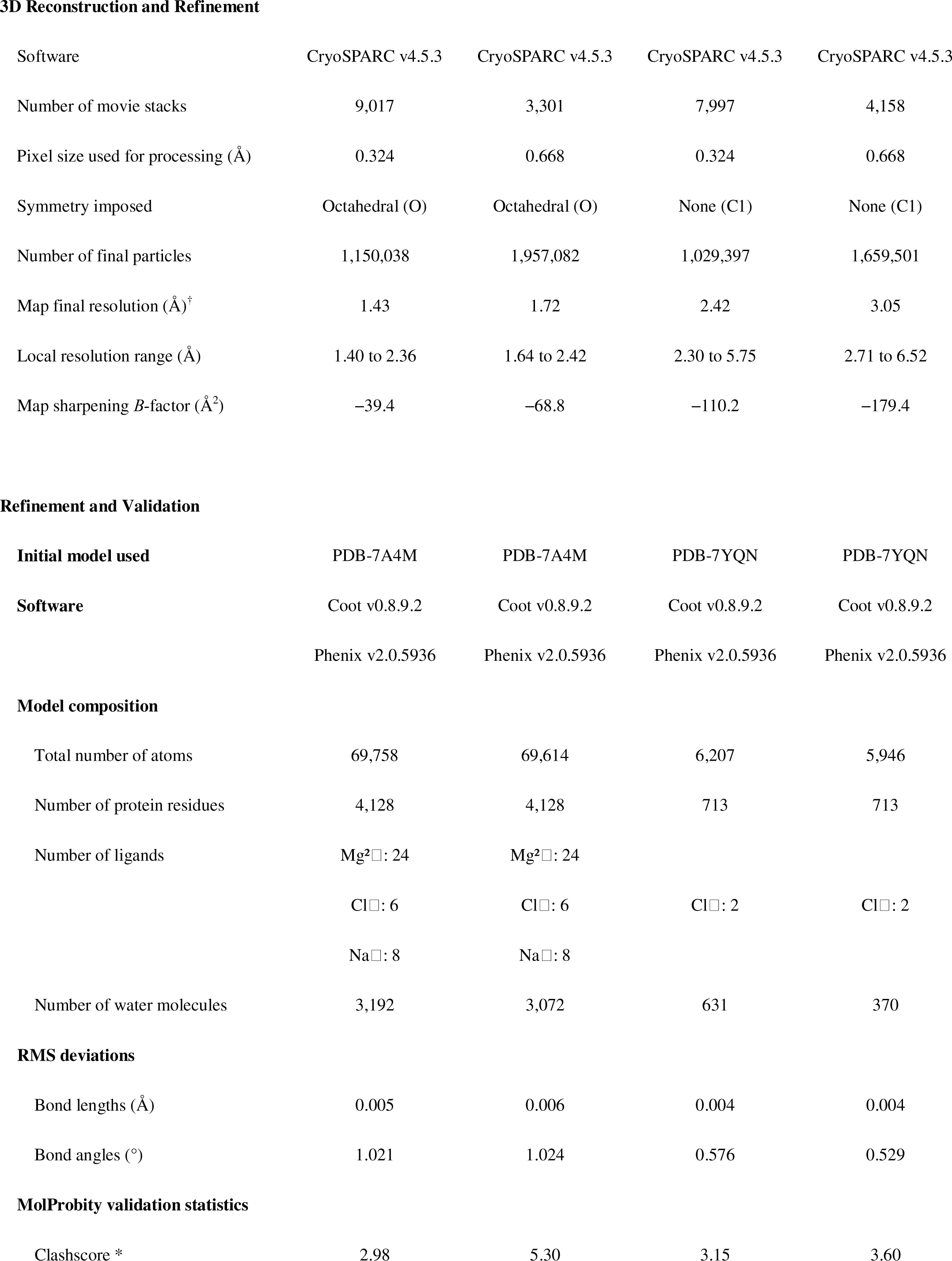
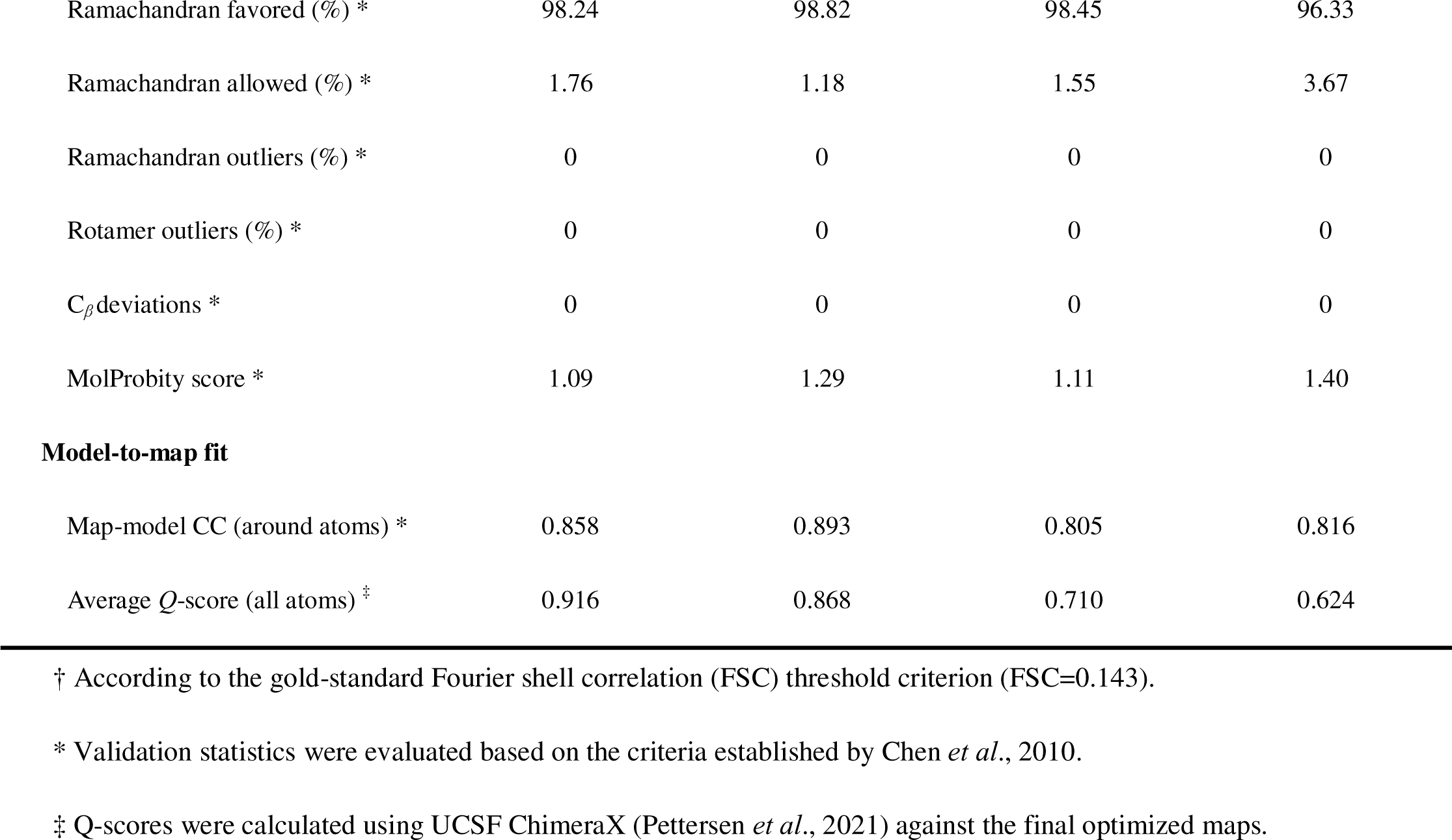
Cryo-EM data collection, 3D reconstruction, and model refinement statistics (Bin1 with per-particle motion correction).

|  | Apoferritin |  | Malate synthase G |  |
| --- | --- | --- | --- | --- |
|  | 130,000x | 64,000x | 130,000x | 64,000x |
|  | (EMD-81736) | (EMD-81737) | (EMD-81738) | (EMD-81739) |
|  | (PDB-28GV) | (PDB-28GY) | (PDB-28GZ) | (PDB-28HA) |
| Data collection |  |  |  |  |
| Microscope | Titan Krios | Titan Krios | Titan Krios | Titan Krios |
| Acceleration voltage (kV) | 300 | 300 | 300 | 300 |
| Spherical aberration, Cs (mm) | 2.7 | 2.7 | 2.7 | 2.7 |
| Detector (Energy filter) | K3 (BioQuantum) | K3 (BioQuantum) | K3 (BioQuantum) | K3 (BioQuantum) |
| Detector mode | Super-resolution | Super-resolution | Super-resolution | Super-resolution |
| Slit width (eV) | 15 | 15 | 15 | 15 |
| Grid type | HexAuFoil | UltrAuFoil | UltrAuFoil | UltrAuFoil |
|  | (R0.29/0.31) | (R1.2/1.3) | (R1.2/1.3) | (R1.2/1.3) |
| Nominal magnification (×) | 130,000 | 64,000 | 130,000 | 64,000 |
| Physical pixel size (Å) | 0.648 | 1.336 | 0.648 | 1.336 |
| Super-resolution pixel size (Å) | 0.324 | 0.668 | 0.324 | 0.668 |
| Spot size | 4 | 3 | 3 | 3 |
| Target defocus range (μm) | −0.20 to −1.20 | −0.80 to −1.60 | −0.80 to −1.60 | −1.00 to −1.80 |
| Actual defocus range (μm) | −0.35 to −2.94 | −0.35 to −3.00 | −0.36 to −3.00 | −0.36 to −2.99 |
| Exposure time (s) | 1.61 | 2.01 | 1.34 | 2.01 |
| Movie frames (no.) | 60 | 50 | 50 | 50 |
| Dose rate (e<sup>−</sup>/pixel/s) | 3.35 | 11.54 | 4.23 | 11.52 |
| Dose per frame (e<sup>−</sup>/Å<sup>2</sup>/frame) | 0.86 | 1.04 | 1.08 | 1.04 |
| Total electron exposure (e<sup>−</sup>/Å<sup>2</sup>) | 51.4 | 52.0 | 54.0 | 51.9 |
| Data collection software | EPU 3.3.1 | EPU 3.6.0 | EPU 3.6.0 | EPU 3.6.0 |
| 3D Reconstruction and Refinement |  |  |  |  |
| Software | CryoSPARC v4.5.3 | CryoSPARC v4.5.3 | CryoSPARC v4.5.3 | CryoSPARC v4.5.3 |
| Number of movie stacks | 9,017 | 3,301 | 7,997 | 4,158 |
| Pixel size used for processing (Å) | 0.324 | 0.668 | 0.324 | 0.668 |
| Symmetry imposed | Octahedral (O) | Octahedral (O) | None (C1) | None (C1) |
| Number of final particles | 1,150,038 | 1,957,082 | 1,029,397 | 1,659,501 |
| Map final resolution (Å)<sup>†</sup> | 1.43 | 1.72 | 2.42 | 3.05 |
| Local resolution range (Å) | 1.40 to 2.36 | 1.64 to 2.42 | 2.30 to 5.75 | 2.71 to 6.52 |
| Map sharpening *B*-factor (Å<sup>2</sup>) | −39.4 | −68.8 | −110.2 | −179.4 |
| Refinement and Validation |  |  |  |  |
| Initial model used | PDB-7A4M | PDB-7A4M | PDB-7YQN | PDB-7YQN |
| Software | Coot v0.8.9.2 | Coot v0.8.9.2 | Coot v0.8.9.2 | Coot v0.8.9.2 |
|  | Phenix v2.0.5936 | Phenix v2.0.5936 | Phenix v2.0.5936 | Phenix v2.0.5936 |
| Model composition |  |  |  |  |
| Total number of atoms | 69,758 | 69,614 | 6,207 | 5,946 |
| Number of protein residues | 4,128 | 4,128 | 713 | 713 |
| Number of ligands | Mg<sup>2+</sup>: 24 | Mg<sup>2+</sup>: 24 |  |  |
|  | Cl<sup>−</sup>: 6 | Cl<sup>−</sup>: 6 | Cl<sup>−</sup>: 2 | Cl<sup>−</sup>: 2 |
|  | Na<sup>+</sup>: 8 | Na<sup>+</sup>: 8 |  |  |
| Number of water molecules | 3,192 | 3,072 | 631 | 370 |
| RMS deviations |  |  |  |  |
| Bond lengths (Å) | 0.005 | 0.006 | 0.004 | 0.004 |
| Bond angles (°) | 1.021 | 1.024 | 0.576 | 0.529 |
| MolProbity validation statistics |  |  |  |  |
| Clashscore * | 2.98 | 5.30 | 3.15 | 3.60 |
| Ramachandran favored (%) * | 98.24 | 98.82 | 98.45 | 96.33 |
| Ramachandran allowed (%) * | 1.76 | 1.18 | 1.55 | 3.67 |
| Ramachandran outliers (%) * | 0 | 0 | 0 | 0 |
| Rotamer outliers (%) * | 0 | 0 | 0 | 0 |
| C<sub>β</sub> deviations * | 0 | 0 | 0 | 0 |
| MolProbity score * | 1.09 | 1.29 | 1.11 | 1.40 |
| Model-to-map fit |  |  |  |  |
| Map-model CC (around atoms) * | 0.858 | 0.893 | 0.805 | 0.816 |
| Average Q-score (all atoms) ‡ | 0.916 | 0.868 | 0.710 | 0.624 |

† According to the gold-standard Fourier shell correlation (FSC) threshold criterion (FSC=0.143).
\* Validation statistics were evaluated based on the criteria established by Chen *et al.*, 2010.
‡ Q-scores were calculated using UCSF ChimeraX (Pettersen *et al.*, 2021) against the final optimized maps.

### 2.4. Common image-processing workflow

All image processing and 3D reconstruction were performed using CryoSPARC (Punjani *et al*., 2017). The Bin1 and Bin2 workflows shared the same initial movie stacks and overall image-processing procedure, with the primary difference being the pixel sampling used for reconstruction. Bin1 retained the native super-resolution sampling, whereas Bin2 used the corresponding physical-pixel sampling.

Initial image processing included patch-based motion correction and CTF estimation using CTFFIND4 (Rohou & Grigorieff, 2015). Particle picking was performed using specimen-specific methods: APO particles were picked by template-based particle picking, whereas MSG particles were picked using the machine-learning-based Topaz software because of the lower image contrast of MSG (Bepler *et al*., 2019).

The picked particles were then subjected to iterative 2D classification, *ab initio* reconstruction, and 3D heterogeneous refinement. Following these steps, the predominant structural populations were selected for further refinement. The resulting APO particles were refined using homogeneous refinement with octahedral (O) symmetry, whereas MSG particles were refined using non-uniform refinement without imposed symmetry (C1). Per-particle motion correction was then applied before the final refinement using the “Reference Based Motion Correction” job in CryoSPARC (Section 2.8). Detailed processing parameters for each dataset are summarized in Table 1, and the corresponding processing workflows are provided in Figs. S2, S3, S7, and S8.

### 2.5. Model building and validation

Previously reported high-resolution structures were used as starting models, including the 1.22 Å cryo-EM structure of APO (PDB: 7A4M; Nakane *et al*., 2020) and the 1.60 Å X-ray structure of MSG (PDB: 7YQN; Ho *et al*., 2023). Initial rigid-body fitting was performed using UCSF Chimera (Yang *et al*., 2012), followed by manual model adjustment in COOT (Emsley *et al*., 2010) and iterative real-space refinement in PHENIX (Afonine *et al*., 2018). Models were manually rebuilt and refined until convergence and subsequently evaluated using the PHENIX comprehensive cryo-EM validation suite.

Four final atomic models were generated from the per-particle motion-corrected Bin1 maps for APO and MSG at both magnifications (130,000× and 64,000×) (Figs. S6 and S11). These models were subsequently used as common references for Q-score analysis across the corresponding sampling and motion-correction conditions.

### 2.6. Reconstruction evaluation

To systematically evaluate the quality of the final reconstructions, we assessed global and local resolution and density-map features as complementary measures of reconstruction quality. Global resolution of each final reconstruction was determined using the gold-standard Fourier shell correlation (FSC) criterion at FSC = 0.143. Local resolution was estimated using CryoSPARC (Figs. S4, S5, S9, and S10). Final density maps were visualized using UCSF Chimera (Yang *et al*., 2012) and inspected for structural features including backbone continuity, side-chain conformations, coordinated ion densities, and ordered water molecules.

### 2.7. Particle-number saturation analysis

The dependence of reconstruction resolution on particle number was evaluated by generating randomly selected particle subsets from each final particle stack. For each condition, particle subsets were generated at predefined increments, starting from 10,000 particles (200 particles for the APO 64,000× Bin2 dataset) up to the full dataset. The final dataset sizes ranged from ∼1.02 million to ∼1.96 million particles across the four APO conditions, and from ∼1.03 million to ∼1.66 million across the four MSG conditions. Each particle subset was independently refined using the same reconstruction strategy as the corresponding full dataset: homogeneous refinement with octahedral (O) symmetry for APO and non-uniform refinement without symmetry constraints (C1) for MSG. Resolution was determined for each reconstruction using the gold-standard FSC = 0.143 criterion (Figs. S12 and S13). Resolution was plotted as a function of particle number for all eight experimental conditions (two specimens × two magnifications × two pixel-sampling strategies), generating the particle-number saturation curves shown in Fig. 5.

### 2.8. Per-particle motion correction analysis

The contribution of per-particle motion correction to reconstruction quality was assessed using the “Reference Based Motion Correction” job in CryoSPARC. For each of the eight experimental conditions, the same particle stack was refined before and after per-particle motion correction using otherwise identical refinement parameters. Global resolution was determined using the gold-standard FSC = 0.143 criterion, and the corresponding resolution changes are summarized in Fig. 6 and Table S1. Differences in resolution before and after correction were used to assess the effect of per-particle motion correction under each magnification and pixel-sampling condition. Q-score analysis was performed separately as described in Section 2.9.

### 2.9. Q-score analysis

To quantitatively assess map quality, map-to-model agreement was evaluated using Q-scores (Pintilie *et al*., 2020) calculated in UCSF ChimeraX (Pettersen *et al*., 2021). For each model, four corresponding maps were evaluated: Bin1 before and after per-particle motion correction and Bin2 before and after per-particle motion correction. The models were rigid-body fitted without additional model refinement, and Q-scores were calculated for each map condition (Table S1). Because the same refined model was used for all four corresponding maps, differences in Q-scores primarily reflected differences in map quality rather than differences in model geometry.

## 3. Results

### 3.1. Experimental design for assessing sampling and signal limitations

To determine when low-magnification super-resolution imaging improves cryo-EM reconstruction, we established a matched benchmarking framework using two structurally distinct specimens. Apoferritin (APO, ∼500 kDa) was selected as a large, highly symmetric, high-contrast specimen, whereas malate synthase G (MSG, ∼80 kDa) represented a smaller, asymmetric, lower-contrast protein. For each specimen, independent datasets were collected at 130,000× and 64,000× nominal magnifications using the same microscope and detector configuration. Each dataset was then processed from the same movie stacks using two parallel workflows: super-resolution processing at the native super-resolution pixel size (Bin1) and physical-pixel processing at the detector pixel size (Bin2). This design allowed the effects of acquisition magnification and image-sampling strategy to be evaluated independently. We used global resolution, density-map features, particle-number saturation, per-particle motion correction, and Q-score analysis to compare the resulting reconstructions. This multi-metric approach was designed to evaluate how sampling bandwidth and recoverable particle signal relate to reconstruction quality under different experimental conditions.

### 3.2. Low-magnification APO reconstruction exhibits sampling-limited behavior

We first established a high-magnification baseline for APO at 130,000× (Fig. 1A). 2D classification of APO produced highly similar class averages under both the Bin1 and Bin2 workflows (∼4.0 Å; Fig. 1C). Global FSC analysis confirmed near-identical final 3D resolutions of 1.43 Å for Bin1 and 1.45 Å for Bin2 (Fig. 1E), with both reconstructions resolving detailed side-chain rotamers, carbonyl oxygen orientations, coordinated ions, and ordered hydration networks (Figs. 2A and S6E). Thus, both workflows produced essentially equivalent reconstructions at 130,000×.

**Figure 1.**
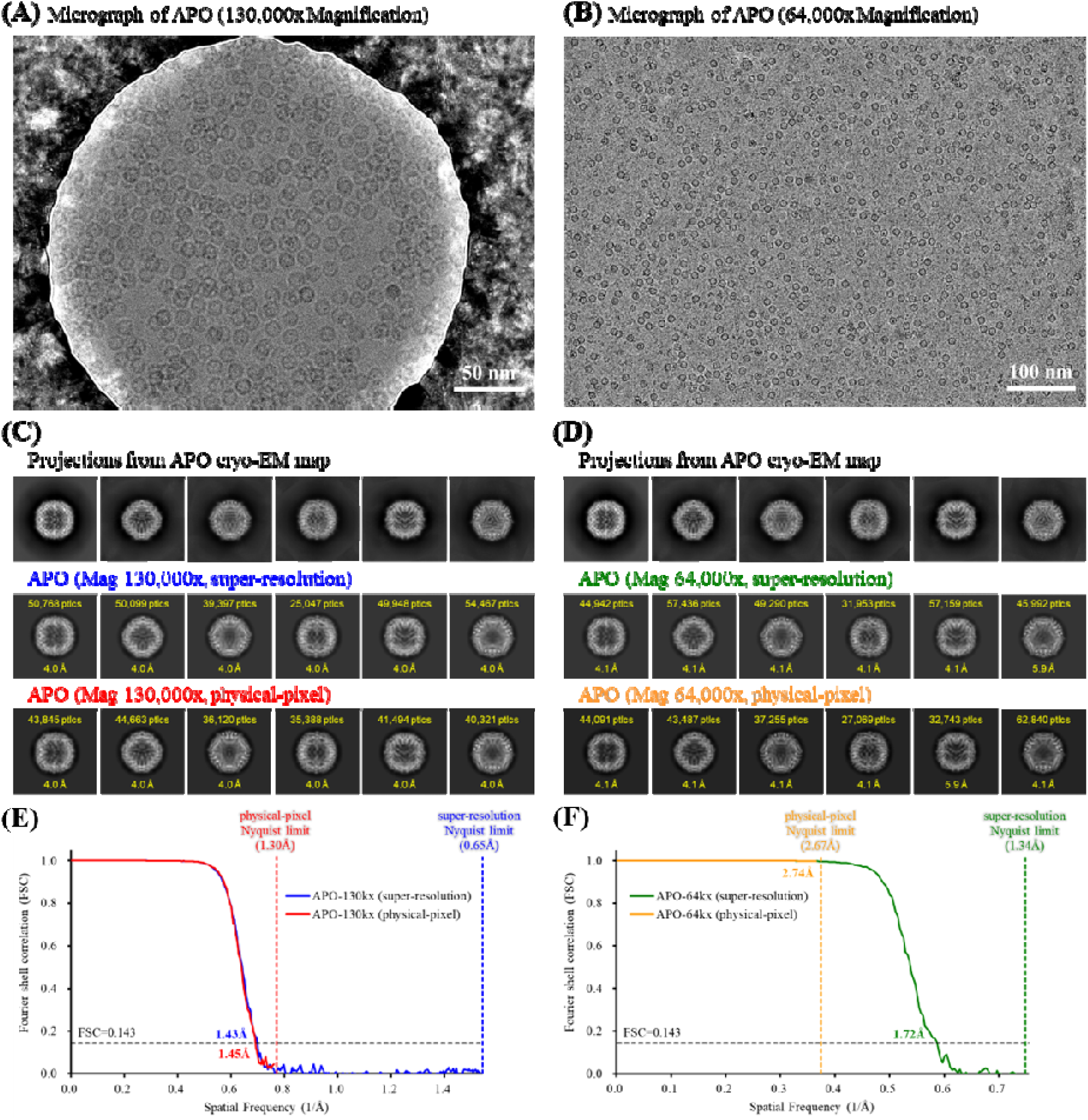
Global reconstruction quality and resolution assessment of apoferritin (APO) at 130,000× and 64,000× magnifications. **(A–B)** Representative cryo-EM micrographs acquired at 130,000× and super-resolution (Bin1) and physical-pixel (Bin2) workflows at 130,000× and 64,000×, respectively, with corresponding projections from the cryo-EM maps shown in the top rows. Particle counts and estimated resolutions are indicated for each class. **(E–F)** Gold-standard FSC curves for the 130,000× and 64,000× datasets, respectively. Global map resolutions were determined using the FSC = 0.143 criterion (horizontal dashed line). Vertical dashed lines indicate the respective physical-pixel and super-resolution Nyquist limits.

**Figure 2.**
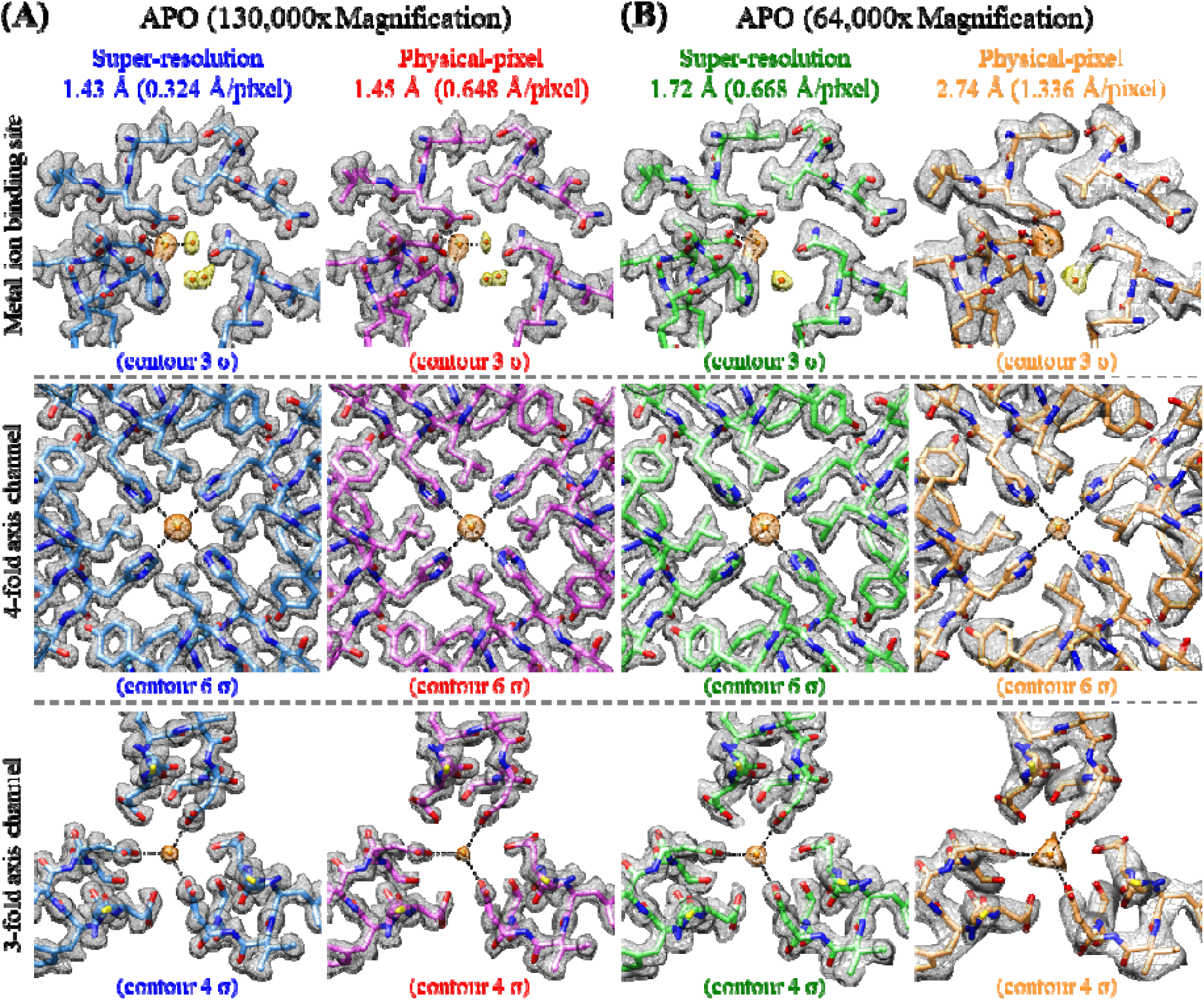
Local density-map interpretability and structural detail preservation of apoferritin (APO) across magnifications and sampling strategies. Representative structural regions are compared between super-resolution (Bin1) and physical-pixel (Bin2) reconstructions obtained at **(A)** 130,000× and **(B)** 64,000× magnifications after per-particle motion correction. The selected regions include a metal ion binding site (top row, contoured at 3 σ), the 4-fold axis channel (middle row, contoured at 6 σ), and the 3-fold axis channel (bottom row, contoured at 4 σ). Global map resolutions and pixel sizes for each condition are annotated at the top of each column. Density maps are rendered as transparent gray meshes overlaid with the corresponding atomic models shown as colored sticks. These contour thresholds were selected to optimally visualize structural features for each reconstruction. Coordinated water molecules (H□O) and other ions are represented as red and orange spheres, respectively.

We next evaluated APO at the lower magnification of 64,000× (Fig. 1B). The nominal field of view increased 4.25-fold (∼98,973 nm² at 130,000× versus ∼420,686 nm² at 64,000×), corresponding to an approximately 4.9-fold difference in observed particle yield (∼120 versus ∼590 particles per micrograph, Fig. 1A and 1B). This difference exceeded the nominal field-of-view scaling alone, likely reflecting differences in the effective imaging area between the two grid types. At 130,000×, the imaging area was partially obstructed by the gold substrate borders of the small-hole HexAuFoil grid (Naydenova *et al*., 2020). Conversely, the larger-hole UltrAuFoil grid used at 64,000× allowed the expanded field of view to be more fully utilized within open ice. Under this low-magnification condition, 2D classification produced highly similar class averages at ∼4.1 Å for both workflows (Fig. 1D), with only minor differences in particle assignment among low-population orientations.

In contrast, the final 3D reconstructions diverged substantially: the Bin2 physical-pixel reconstruction plateaued at 2.74 Å, close to the physical Nyquist limit of 2.67 Å, whereas the Bin1 super-resolution reconstruction reached 1.72 Å (Fig. 1F), extending well beyond the physical Nyquist limit. This improvement indicates that super-resolution imaging can recover high-spatial-frequency information beyond the physical Nyquist limit, consistent with previous studies (Feathers *et al*., 2021; Sheng *et al*., 2022). Visually, the Bin1 map retained detailed side-chain features and ion coordination sites that were less well defined in the Bin2 map (Figs. 2B and S6F). Together, these findings indicate that, under the present conditions, the 64,000× APO reconstruction exhibited sampling-limited behavior.

### 3.3. Low-magnification MSG reconstruction exhibits predominantly signal-limited behavior

We next examined MSG under the same acquisition and processing framework (Fig. 3A and 3B). At 130,000×, both workflows for MSG produced high-quality 2D class averages resolving the predominant orientation classes at ∼3.5–4.5 Å (Fig. 3C). Although minor orientation classes showed slight differences in particle assignment and estimated resolution between the workflows, these local variations did not affect the final 3D reconstruction. FSC analysis confirmed near-identical final resolutions of 2.42 Å for Bin1 and 2.49 Å for Bin2 (Fig. 3E), with both maps resolving the protein backbone, side-chains, and coordinated ion densities (Fig. 4A).

**Figure 3.**
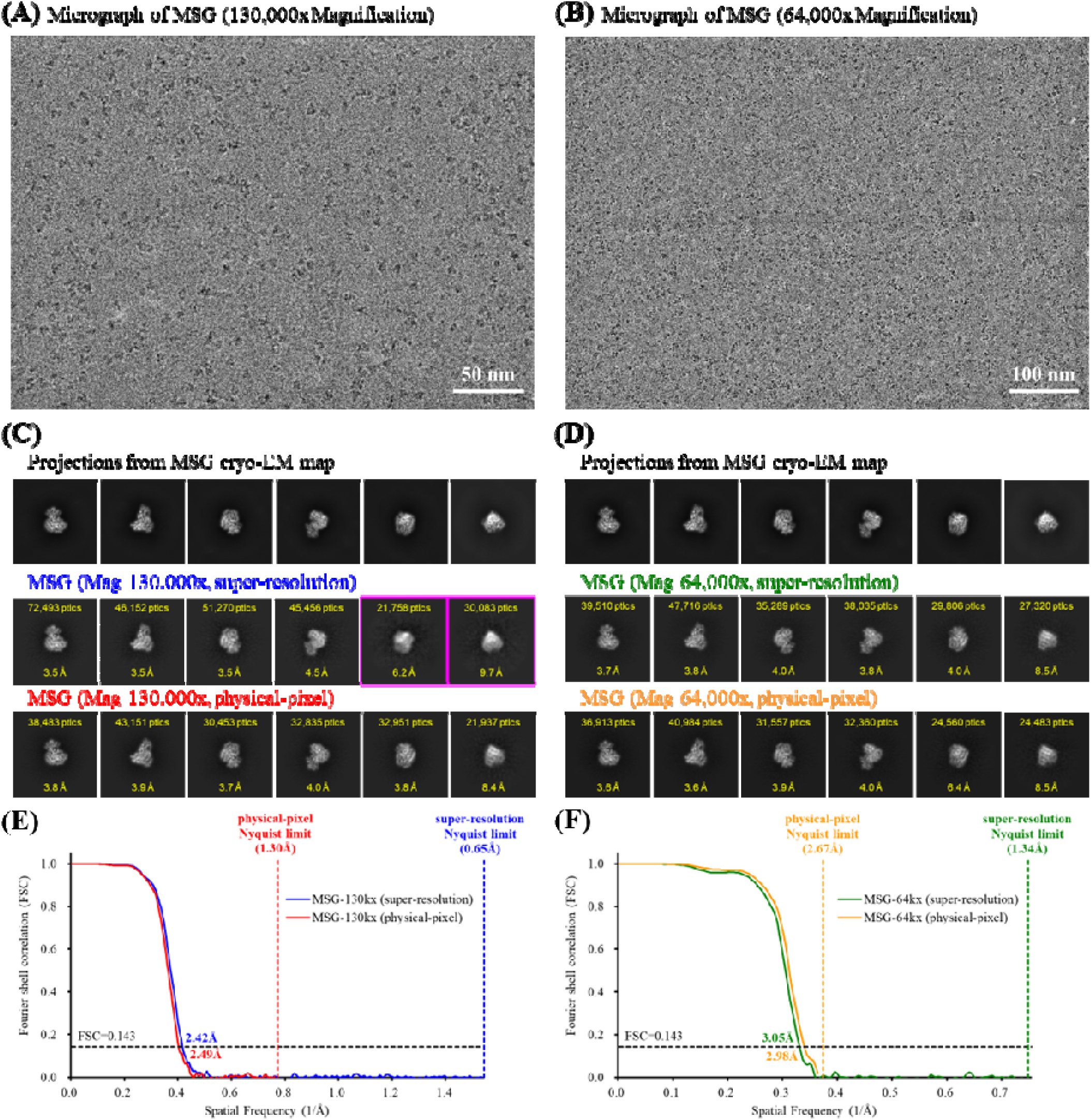
Global reconstruction quality and resolution assessment of malate synthase G (MSG) at 130,000× and 64,000× magnifications. **(A–B)** Representative cryo-EM micrographs acquired at 130,000× and 64,000×, respectively; scale bars, 50 nm **(A)** and 100 nm **(B)**. **(C–D)** Representative 2D class averages from super-resolution (Bin1) and physical-pixel (Bin2) workflows at 130,000× and 64,000×, respectively, with corresponding projections from the cryo-EM maps shown in the top rows. Particle counts and estimated resolutions are indicated for each class. Magenta boxes highlight representative poorly resolved 2D classes with limited structural features. **(E–F)** Gold-standard FSC curves for the 130,000× and 64,000× datasets, respectively. Global map resolutions were determined using the FSC = 0.143 criterion (horizontal dashed line). Vertical dashed lines indicate the respective physical-pixel and super-resolution Nyquist limits.

**Figure 4.**
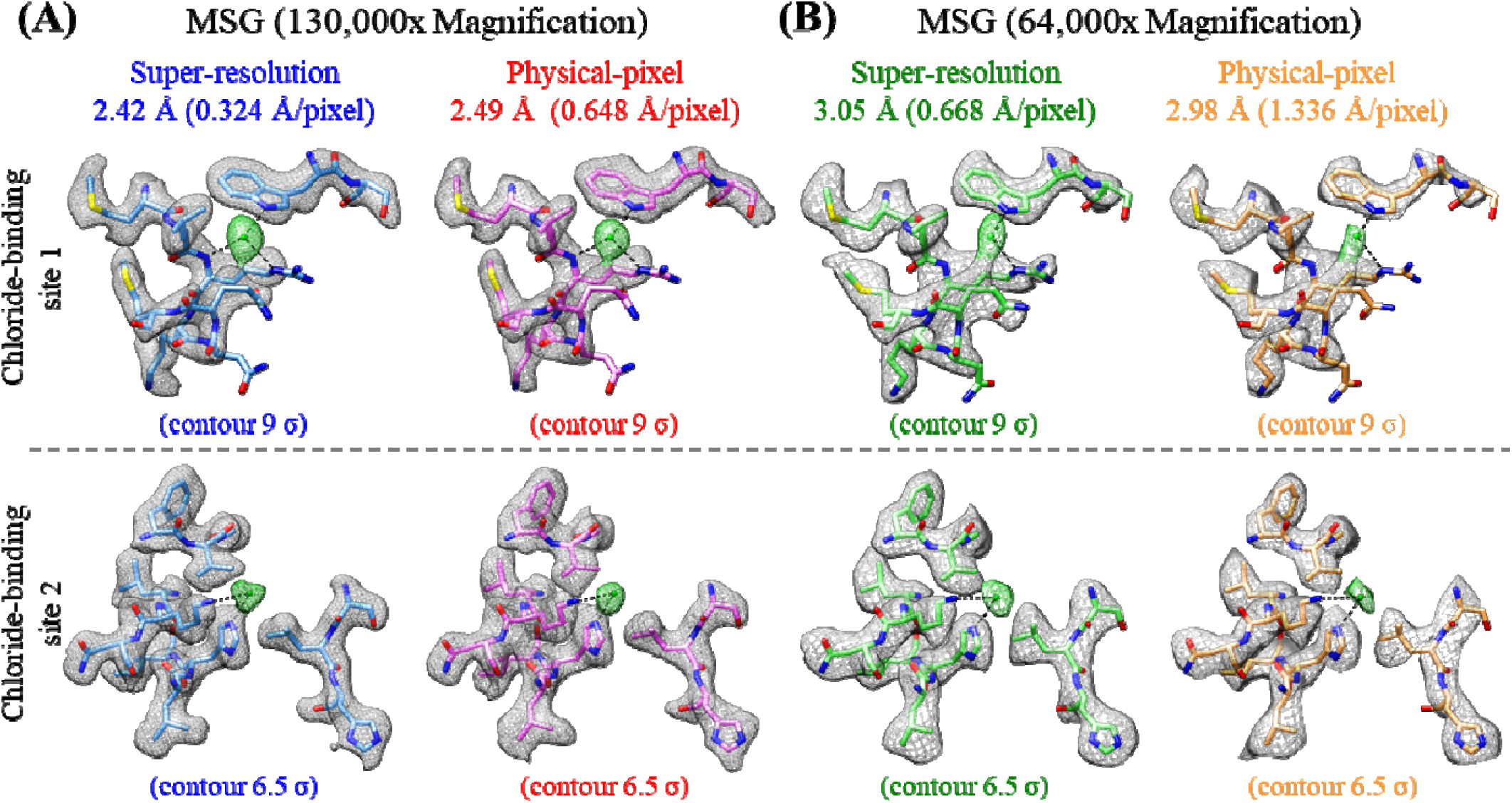
Local density-map interpretability and structural detail preservation of malate synthase G (MSG) across magnifications and sampling strategies. Representative structural regions are compared for super-resolution (Bin1) and physical-pixel (Bin2) reconstructions at **(A)** 130,000× and **(B)** 64,000× magnifications. The selected regions include chloride-binding site 1 (top row, contoured at 9 σ) and chloride-binding site 2 (bottom row, contoured at 6.5 σ). Global map resolutions and pixel sizes for each condition are annotated at the top of each column. Density maps are rendered as transparent gray meshes overlaid with the corresponding atomic models shown as colored sticks. These contour thresholds were selected to optimally visualize structural features for each reconstruction. Chloride ions (Cl□) are represented as green spheres.

At 64,000×, the 4.25-fold field-of-view expansion yielded an approximately 3.1-fold increase in practical particle throughput (∼130 versus ∼400 particles per micrograph; Fig. 3A and 3B). 2D classification again produced visually comparable class averages between the workflows (∼3.6–4.0 Å; Fig. 3D). However, final 3D refinement converged to similar resolutions of 3.05 Å for Bin1 and 2.98 Å for Bin2 (Fig. 3F). Both resolutions remained substantially coarser than the physical Nyquist limit of 2.67 Å, indicating that the available physical-pixel sampling bandwidth was not the dominant limitation under this condition. Although the backbone density remained continuous under both conditions, coordinated ion densities and other local density features were slightly better preserved in the Bin1 map (Fig. 4B). Together, these findings indicate that, under the present conditions, the 64,000× MSG reconstruction exhibited predominantly signal-limited behavior.

### 3.4. Complementary analyses support the distinct reconstruction behaviors

Because the 130,000× datasets showed near-identical Bin1 and Bin2 reconstructions for both specimens, subsequent analyses focused on the 64,000× datasets, where the two specimens exhibited markedly different responses to the two sampling strategies. Three complementary analyses were employed to examine how particle number, per-particle motion correction, and local map quality related to these observed differences.

First, particle-number saturation analysis evaluated reconstruction resolution as a function of dataset size (Fig. 5). For APO at 64,000×, the Bin2 workflow rapidly plateaued at 2.74 Å near the physical Nyquist boundary, whereas the Bin1 workflow continued to improve steadily with increasing particle numbers, extending well beyond this limit. In contrast, both 64,000× MSG workflows exhibited highly similar trajectories that plateaued well below the physical Nyquist limit, showing progressively diminishing resolution gains as particle numbers increased. These distinct trajectories were consistent with the contrasting sampling-limited behavior of APO and predominantly signal-limited behavior of MSG.

**Figure 5.**
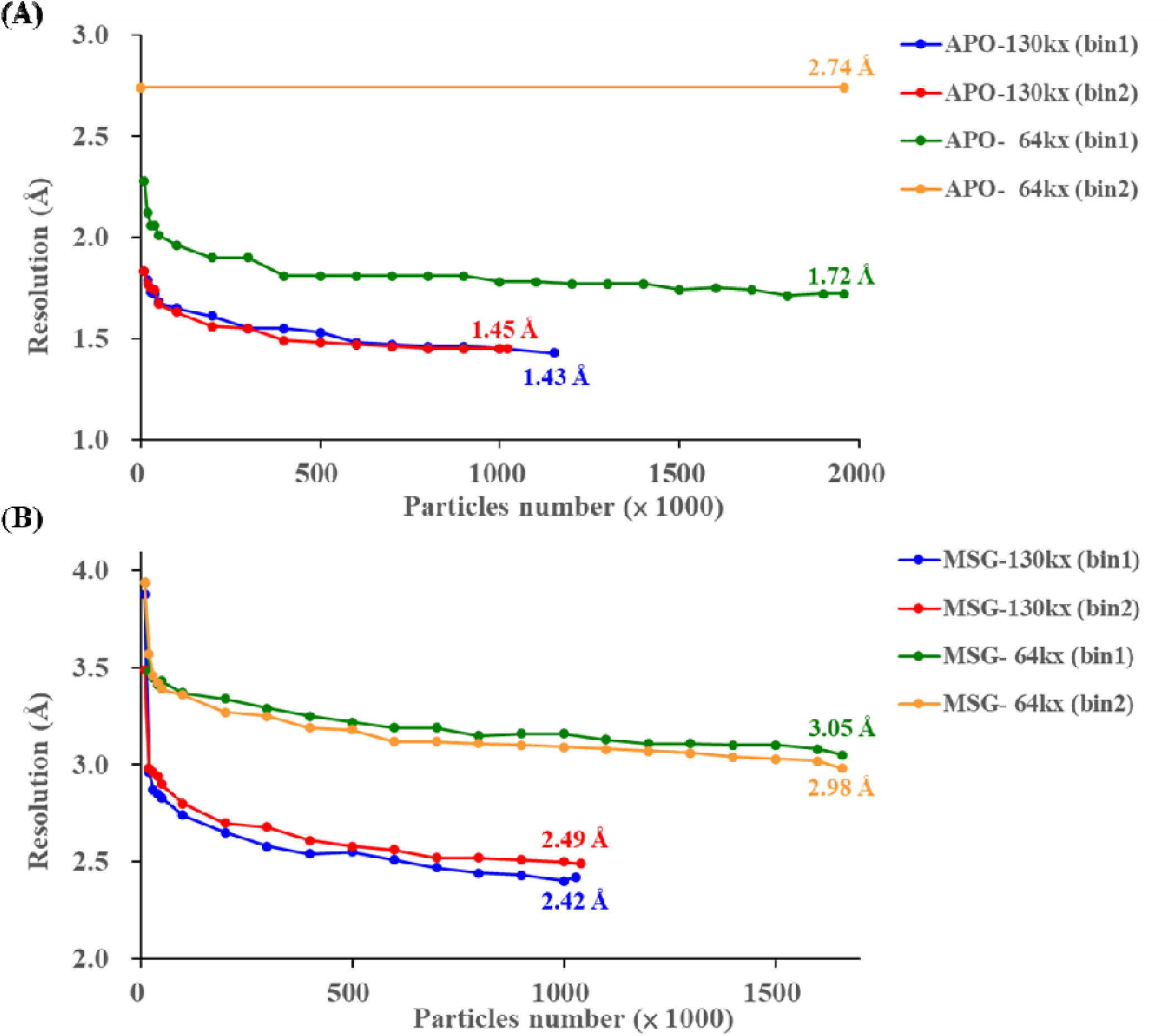
Particle-number dependence and resolution saturation profiles of apoferritin (APO) and malate synthase G (MSG) reconstructions. Global 3D reconstruction resolutions are plotted as a function of randomly subsampled particle numbers for super-resolution (Bin1) and physical-pixel (Bin2) workflows at 130,000× and 64,000× magnifications. Sampling trajectories are shown for **(A)** the larger, symmetric benchmark APO and **(B)** the smaller, asymmetric target MSG. Each curve represents the resolution achieved from an increasing subset of the corresponding final particle stack. Global resolutions were determined using the gold-standard FSC = 0.143 criterion. This analysis was used to assess whether reconstruction performance was limited by detector sampling or by available particle signal.

Second, the effect of per-particle motion correction on reconstruction quality was assessed by comparing maps before and after its application (Fig. 6). This correction accounts for residual particle-specific motion and can improve the recovery of high-resolution structural details (Zivanov *et al*., 2019). For APO at 64,000×, per-particle motion correction improved the global resolution of the Bin1 reconstruction from 1.86 Å to 1.72 Å, whereas the Bin2 reconstruction remained at 2.74 Å, close to the physical Nyquist limit. Consistent with this global resolution improvement, local-resolution analysis revealed enhanced resolution in several regions of the APO Bin1 map following correction (Figs. S4 and S5). This differential response is consistent with the interpretation that improved particle-level motion estimation can enhance reconstruction quality when additional high-frequency information remains recoverable and sufficient sampling bandwidth is available to represent it. In contrast, the Bin2 reconstruction remained unchanged at 2.74 Å, consistent with the physical Nyquist limit representing the dominant constraint on further resolution improvement under this condition. For MSG, per-particle motion correction produced only modest changes in both global and local resolutions, indicating limited additional improvement under the reconstruction conditions examined here.

**Figure 6.**
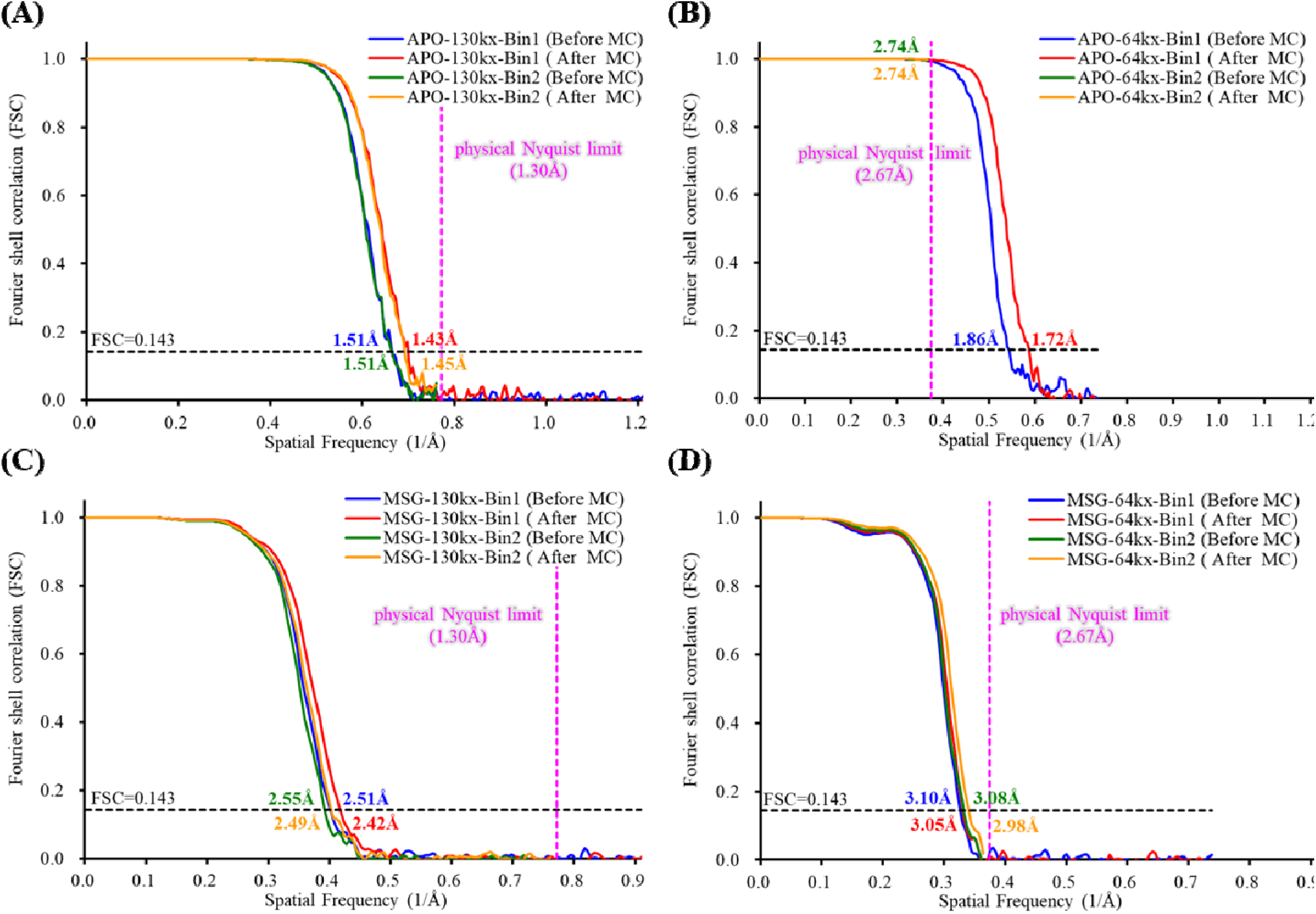
Effect of per-particle motion correction on reconstruction resolution of apoferritin (APO) and malate synthase G (MSG) across magnifications and sampling conditions. Gold-standard FSC curves before and after per-particle motion correction (MC) are systematically compared for reconstructions under different imaging and sampling conditions: **(A)** APO at 130,000×, **(B)** APO at 64,000×, **(C)** MSG at 130,000×, and **(D)** MSG at 64,000× magnifications. Global map resolutions were determined using the gold-standard FSC = 0.143 criterion (horizontal dashed line). Vertical dashed lines indicate the corresponding physical-pixel Nyquist limits.

Third, quantitative Q-score analysis provided an independent assessment of local map quality (Pintilie *et al*., 2020; Table 1; Table S1). For APO at 64,000×, the Q-scores differed substantially between Bin1 and Bin2 (0.868 versus 0.648), consistent with the pronounced differences in local density features observed between the two maps (Figs. 2B and S6F). For MSG at 64,000×, the corresponding difference was much smaller (0.624 versus 0.598), consistent with the broadly similar map quality observed between the two workflows, while still indicating modest local differences despite their similar global resolutions (Figs. 4B and S11F).

Taken together, these analyses provided complementary evidence for the distinct responses of APO and MSG to changes in sampling strategy. Particle-number saturation showed the clearest divergence between the two specimens, while per-particle motion correction and Q-score analysis provided complementary evidence for high-resolution information recovery and local map quality, respectively.

## 4. Discussion

### 4.1. Sampling- and signal-limited regimes explain the contrasting responses of APO and MSG

The central finding of this study is that the benefit of low-magnification super-resolution imaging is strongly specimen-dependent and can differ substantially among molecular targets. Under matched acquisition and processing conditions, APO showed a pronounced improvement when super-resolution sampling was retained, whereas MSG showed no corresponding benefit. At 64,000×, physical-pixel processing (Bin2) yielded a marginally higher global resolution than super-resolution processing (Bin1), opposite to the response observed for APO.

This divergence indicates that the benefit of super-resolution depends not simply on pixel size, but on whether recoverable structural information extends beyond the sampling bandwidth represented by the physical-pixel image. For APO, sufficient high-frequency signal remained available beyond the physical-pixel sampling rate, such that the physical Nyquist limit became a dominant bottleneck. Retaining super-resolution sampling therefore allowed this high-frequency information to be represented in the reconstruction, consistent with previous demonstrations of information recovery beyond the conventional detector sampling boundary (Li *et al*., 2013; Feathers *et al*., 2021; Sheng *et al*., 2022). This behavior is consistent with a sampling-limited regime.

For MSG, by contrast, the available high-frequency information was primarily constrained by recoverable particle signal rather than by detector sampling. Smaller, lower-contrast particles provide less alignment-informative signal per particle, and reduced image contrast can compromise particle picking, orientation assignment, and high-resolution refinement, making alignment increasingly sensitive to noise (Herzik *et al*., 2017; Wu & Lander, 2020; Wentinck *et al*., 2022). Under these conditions, the expanded sampling bandwidth provided by super-resolution does not necessarily translate into additional recoverable structural information.

Moreover, retaining finer sampling in the super-resolution workflow does not necessarily translate into improved alignment or refinement performance, which may contribute to the slightly higher global resolution obtained with physical-pixel processing for MSG at 64,000×. Additional factors, including beam-induced motion and radiation damage, may further constrain high-resolution information recovery in this regime (Scheres, 2014; Baker & Rubinstein, 2010). This behavior is therefore consistent with a signal-limited regime.

These two regimes should not be regarded as fixed intrinsic properties of specific specimens, but rather as descriptions of the dominant limitation under a particular combination of specimen properties, imaging conditions, particle number, and processing strategy. A given specimen may therefore shift between regimes as experimental conditions change. Nevertheless, the present comparison demonstrates that specimens differing in molecular size, symmetry, and image contrast can respond very differently to the same increase in sampling bandwidth.

The key question is therefore not simply whether super-resolution extends the physical Nyquist limit, but whether sufficient recoverable structural information remains available to exploit the extended sampling bandwidth. This distinction provides a conceptual framework for understanding why increasing sampling density can produce substantial improvements for some specimens while providing diminishing returns for others.

### 4.2. Three complementary signatures identify the dominant reconstruction limitation

Because the dominant limitation of a reconstruction is not always discernible from global resolution alone, we evaluated three complementary signatures that provide independent perspectives on the factors limiting reconstruction quality (Figs. 5 and 6; Table 1 and Table S1). Particle-number saturation, per-particle motion correction, and Q-score analysis probe different aspects of information recovery and together provide a more informative assessment than any single metric alone.

Particle-number saturation provides an indication of whether additional particles continue to contribute recoverable high-frequency information. Continued improvement in resolution with increasing particle number suggests that additional information remains available for refinement, whereas early saturation indicates diminishing returns under the current reconstruction conditions. In the present study, the distinct saturation behavior of APO and MSG was therefore consistent with their different responses to sampling bandwidth (Fig. 5). However, saturation behavior alone does not uniquely identify whether sampling or particle signal is responsible for the observed plateau. Its diagnostic value is therefore strongest when interpreted together with the other measures.

Per-particle motion correction offers a complementary mechanistic perspective. Residual particle-specific motion can blur high-resolution information, and improved estimation of particle trajectories can therefore enhance reconstruction quality (Li *et al*., 2013; Zivanov *et al*., 2019). However, the benefit of this correction depends on whether particle-specific information can be reliably estimated and whether the recovered high-frequency information can be represented within the available sampling bandwidth. The different responses observed between APO and MSG therefore suggest that the benefit of motion correction depends on both the quality of the particle signal available for motion estimation and the sampling bandwidth available to represent the resulting high-frequency information (Fig. 6).

Q-score analysis introduces an additional measure of local map quality that can reveal differences not fully captured by global FSC resolution. This is particularly important when two reconstructions achieve similar global resolutions but differ in the quality of local structural features. In the present study, the MSG super-resolution reconstruction showed modestly improved local density features and higher Q-scores despite its marginally lower global resolution (Fig. 4B; Table S1). Thus, relying on global resolution alone could underestimate local structural differences between the two sampling strategies.

Taken together, these analyses should be viewed as complementary diagnostic signatures rather than independent classification criteria. Particle-number saturation indicates whether additional data continue to yield measurable gains, per-particle motion correction probes the recoverability of particle-specific high-resolution information, and Q-scores provide complementary evidence for local map quality. Their combined behavior provides a practical basis for assessing the dominant limitation under a given reconstruction condition.

This multi-metric perspective is important because sampling and signal limitations may contribute simultaneously to reconstruction quality. The distinction between sampling-limited and signal-limited behavior should therefore not be interpreted as a binary classification, but rather as a means of identifying the dominant bottleneck under a given experimental condition. This distinction has practical implications for optimization: when sampling is dominant, increasing sampling bandwidth may enable recovery of additional high-frequency information, whereas when particle signal is dominant, improving signal quality and alignment accuracy is likely to be more effective.

### 4.3. Translating regime diagnosis into acquisition strategy

The distinction between sampling- and signal-limited reconstruction has direct practical implications for cryo-EM data acquisition. Lower magnification increases the field of view and can substantially increase the number of particles recorded per micrograph, but it also results in coarser physical sampling. Whether super-resolution acquisition can compensate for this loss of sampling bandwidth depends on the specimen and the amount of recoverable high-frequency information.

For specimens that retain sufficient high-frequency signal, as exemplified by APO under the conditions examined here, low-magnification super-resolution imaging can provide an effective means of increasing particle throughput while retaining access to information beyond the physical Nyquist limit. This strategy may be particularly advantageous when particle yield is a major practical constraint and the specimen contains sufficient high-frequency information to benefit from the additional sampling bandwidth.

However, when the highest achievable resolution is the primary objective, high-magnification acquisition remains advantageous. Higher magnification provides finer physical sampling without relying on information recovery beyond the physical-pixel Nyquist limit and may provide more favorable conditions for precise particle alignment and high-resolution refinement. Low-magnification super-resolution should therefore not be considered a universal replacement for high-magnification acquisition, but rather an alternative strategy when increased particle throughput provides a meaningful experimental advantage.

For specimens resembling MSG under the conditions examined here, increasing sampling density alone may provide diminishing returns in global resolution. In such cases, optimization should instead focus on factors that enhance the recoverability of particle signal and alignment accuracy, including specimen preparation, image contrast, ice conditions, beam-induced motion, particle distribution, and other sources of image degradation. These improvements may provide greater benefit than expanding sampling bandwidth alone.

Importantly, similar global resolutions do not necessarily imply identical map quality. The MSG results demonstrate that super-resolution processing can provide modest improvements in local density features and Q-scores even when the global FSC resolution is slightly inferior (Fig. 4B; Table S1). Acquisition strategies should therefore be evaluated using multiple measures of reconstruction quality rather than relying exclusively on a single FSC-derived resolution value. Moreover, low-magnification acquisition need not be excluded for smaller targets when throughput or specimen availability is a priority, provided that structurally interpretable resolution remains achievable. Previous studies have demonstrated near-3 Å cryo-EM reconstructions of sub-100-kDa proteins using conventional imaging approaches (Herzik *et al*., 2019).

As a practical guideline, a preliminary pilot acquisition can be used to compare physical-pixel and super-resolution processing before committing to large-scale data collection. If super-resolution continues to provide resolution gains with increasing particle number while physical-pixel processing approaches the physical Nyquist boundary, the dataset is more likely to benefit from the additional sampling bandwidth. Conversely, if the two sampling modes converge at similar resolutions well below the physical Nyquist limit and exhibit comparable saturation behavior, further optimization may be more effectively directed toward improving recoverable particle signal, image quality, or alignment accuracy. The relationships among specimen characteristics, expected reconstruction regime, and acquisition strategy are summarized in Table S2.

### 4.4. Limitations and future directions

Several limitations should be considered when interpreting these results. First, APO and MSG differ simultaneously in molecular size, symmetry, and image contrast, and the present comparison does not isolate the individual contribution of each factor. In particular, the octahedral (O) symmetry of APO provides extensive symmetry-related averaging that is absent in the asymmetric (C1) reconstruction of MSG. This symmetry-related averaging can significantly enhance the effective reconstruction signal and may contribute directly to the robust high-resolution response observed for APO. Consequently, molecular weight alone should not be regarded as a sufficient predictor of whether a specimen will exhibit sampling-limited or signal-limited behavior.

Second, the present results were obtained using a specific microscope, detector, acquisition configuration, and image-processing workflow. The relative importance of sampling bandwidth and recoverable particle signal may change with detector performance, magnification, electron dose, exposure conditions, specimen preparation, or advances in image-processing methods. In addition, the computational, storage, and data-transfer costs associated with super-resolution acquisition were not explicitly evaluated and may become important considerations for large-scale data collection.

Third, the sampling-limited and signal-limited regimes should be regarded as conceptual endpoints of a continuum rather than as mutually exclusive categories. Individual datasets may occupy intermediate conditions in which both sampling bandwidth and recoverable particle signal contribute substantially to the final reconstruction quality. The dominant limitation may also shift as specimen preparation, particle throughput, acquisition conditions, or image-processing strategies are optimized.

Future studies using systematically selected specimens that independently vary molecular size, symmetry, and image contrast will be important for disentangling the contributions of these factors. For example, comparisons of proteins with similar molecular weights but different symmetry classes, or identical symmetry but different molecular sizes, could help determine the relative contributions of symmetry averaging and molecular size to reconstruction behavior. Extending such comparisons to membrane proteins, flexible complexes, and other low-contrast or heterogeneous specimens will further test the generality of the proposed framework. Systematic benchmarking across different direct electron detectors and acquisition conditions may also help establish quantitative criteria for predicting when low-magnification super-resolution is likely to provide a practical advantage.

Despite these limitations, the present results support a specimen-aware framework in which the benefit of low-magnification super-resolution depends on whether reconstruction is primarily constrained by sampling bandwidth or recoverable particle signal. This framework provides a basis for interpreting specimen-dependent responses and for selecting acquisition and processing strategies according to the dominant limitations under a given experimental condition.

## 5. Conclusion

The contrasting sampling-limited and signal-limited behaviors observed for APO and MSG demonstrate that increased sampling bandwidth does not necessarily translate into improved resolution. Rather, the practical value of super-resolution depends on whether recoverable high-frequency information extends beyond the frequency range represented by physical-pixel sampling. For sampling-limited reconstructions, low-magnification super-resolution imaging can increase particle throughput while retaining access to higher-resolution information beyond the physical Nyquist limit. In contrast, when reconstruction quality is primarily constrained by recoverable particle signal, increasing sampling bandwidth alone provides limited gains in global resolution. These findings support a specimen-aware approach to cryo-EM data acquisition, in which magnification and sampling strategies are selected according to the dominant limitation of the reconstruction, providing a practical framework for balancing resolution, particle throughput, and data-collection efficiency.

## Acknowledgements

Cryo-EM data were collected at the Academia Sinica Cryo-EM Facility (ASCEM), supported by the Academia Sinica Core Facility and Innovative Instrument Project (Grant No. AS-CFII-111-210). We thank the staff of the Academia Sinica Cryo-EM Center for their technical support and instrument maintenance. We also acknowledge the Academia Sinica Grid-computing Center (ASGC) for providing distributed cloud computing resources used for image processing, three-dimensional reconstruction, and structural refinement. K.P.W. thanks supported from AS grants AS-IV-115-L0 and AS-IAIA-114-L02 and National Science Technology Council (NSTC) (115-2113-M-001-004-).

## Data availability

The atomic coordinates and corresponding cryo-EM density maps have been deposited in the Protein Data Bank (PDB) and the Electron Microscopy Data Bank (EMDB) under the following accession codes: PDB 28GV and EMD-81736 for the apoferritin (APO) 130,000× dataset; PDB 28GY and EMD-81737 for the APO 64,000× dataset; PDB 28GZ and EMD-81738 for the malate synthase G (MSG) 130,000× dataset; and PDB 28HA and EMD-81739 for the MSG 64,000× dataset. All alternative density maps generated under different processing conditions—including super-resolution (Bin1) and physical-pixel (Bin2) reconstructions, with and without per-particle motion correction—have been deposited as associated maps within the corresponding EMDB entries and are publicly available.

Conceptualization: C.H., Y.C.

Investigation: C.H., K.P., Y.C.

Supervision and funding acquisition: Y.C.

Writing – original draft, review & editing: C.H., K.P., Y.C.

## Notes

### Competing Interest Statement

The authors have declared no competing interest.

