## Supplemental Fig for "Specimen-Dependent Sampling and Signal Limitations Govern the Effectiveness of Low-Magnification Super-Resolution in Cryo-EM"

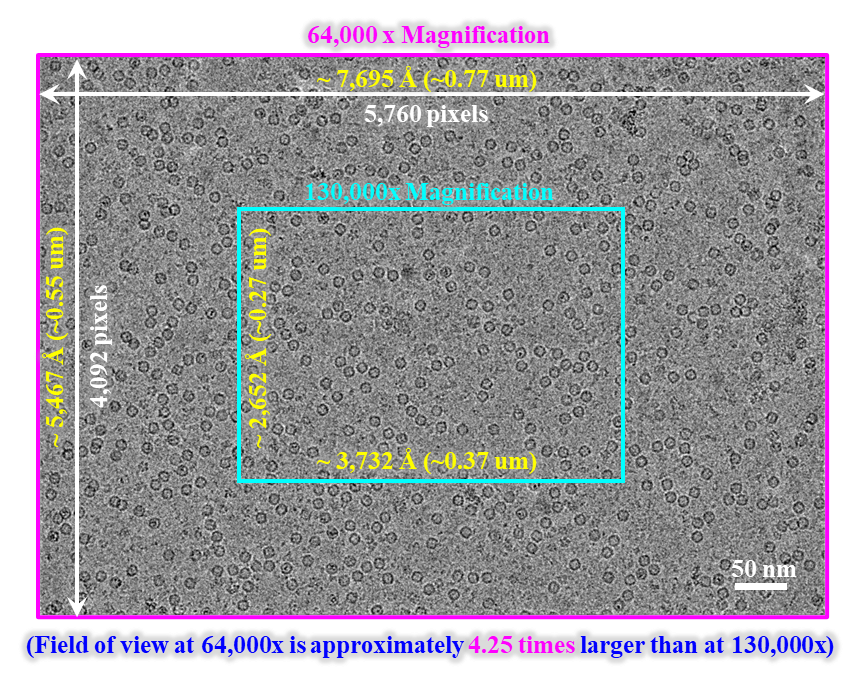

**Fig. S1. Comparison of imaging area at 64,000× and 130,000× magnifications.** A representative cryo-EM micrograph of apoferritin (APO) particles illustrates the difference in imaging area between the two magnifications used in this study. The full micrograph (outlined in pink) corresponds to 64,000× magnification, covering a field of view of ~7,695 × ~5,467 Å (~0.77 × ~0.55 μm) across 5,760 × 4,092 pixels. The cyan box indicates the equivalent field of view at 130,000× magnification, which covers ~3,732 × ~2,652 Å (~0.37 × ~0.27 μm). The imaging area at 64,000× is approximately 4.25-fold larger than that at 130,000×. Individual APO particles are visible as dark, ring-like features distributed across the micrograph. Scale bar, 50 nm.

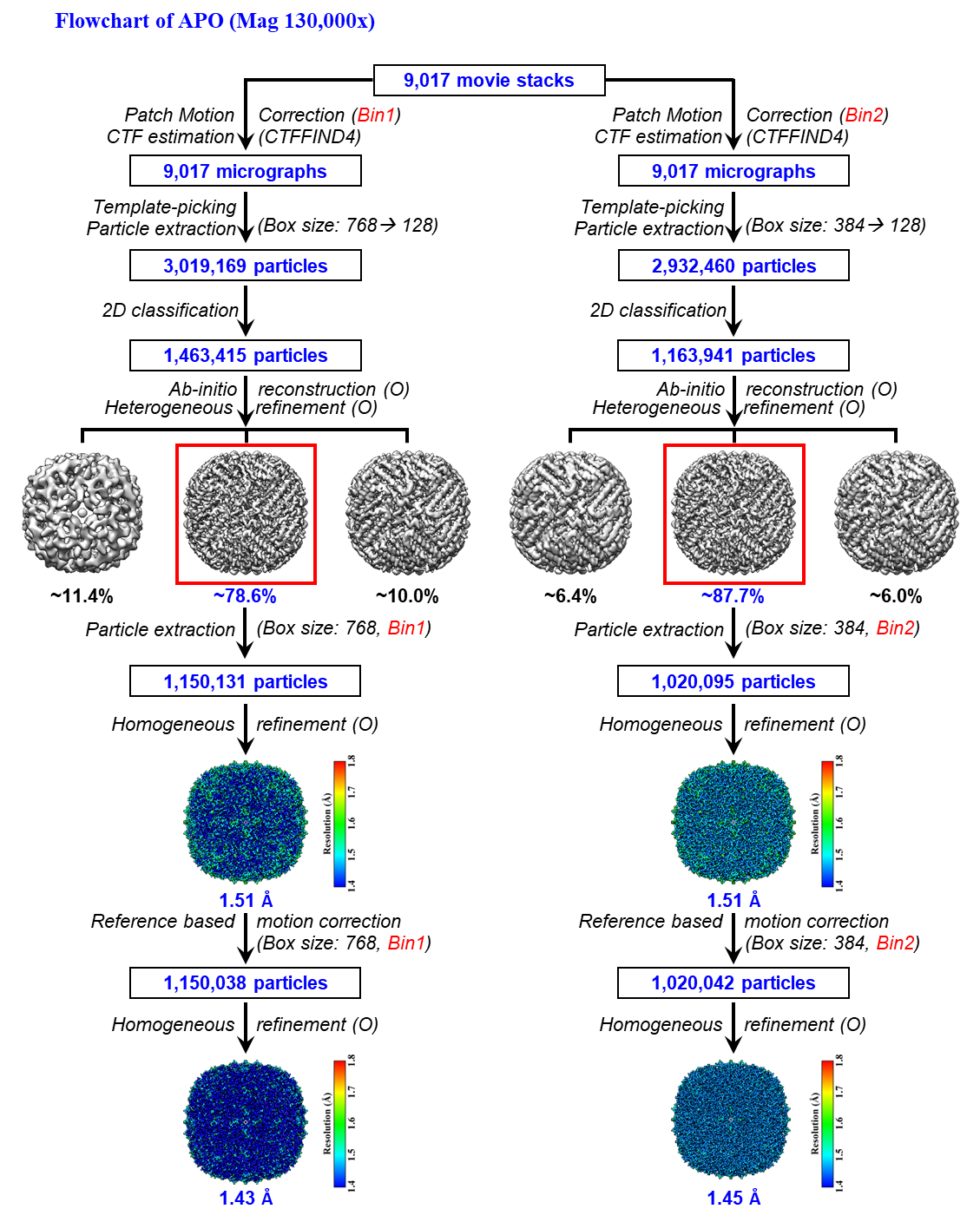

**Figure S2. Cryo-EM data processing workflow for apoferritin (APO) at 130,000× magnification.** The flowchart illustrates the parallel processing strategies derived from the same 9,017 movie stacks using super-resolution (Bin1, left) and physical-pixel (Bin2, right) workflows. Template-based particle picking initially yielded 3,019,169 (Bin1) and 2,932,460 (Bin2) particles. Following 2D classification, 1,463,415 (Bin1) and 1,163,941 (Bin2) particles were selected for *ab-initio* reconstruction and heterogeneous refinement. The predominant 3D class (outlined in red) accounted for ~78.6% (Bin1) and ~87.7% (Bin2) of the particles, resulting in 1,150,131 and 1,020,095 particles after extraction, respectively. Prior to reference-based motion correction, homogeneous refinement yielded an identical global resolution of 1.51 Å for both strategies. Following reference-based motion correction, the final reconstructions using 1,150,038 (Bin1) and 1,020,042 (Bin2) particles achieved resolutions of 1.43 Å and 1.45 Å, respectively. Each local-resolution map (contour at 4 σ) features a color scale on the right and the global resolution noted below.

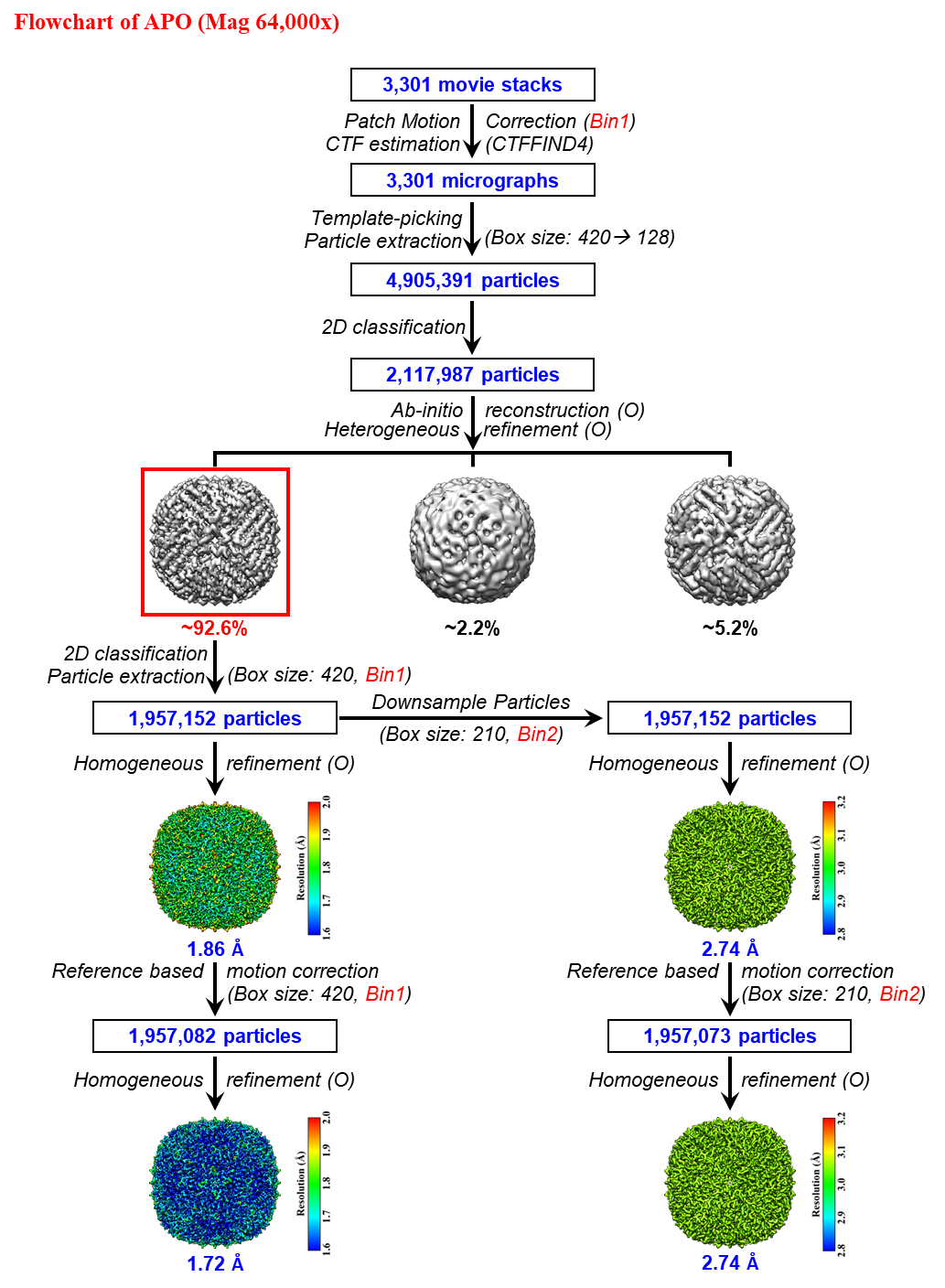

**Figure S3. Cryo-EM data processing workflow for apoferritin (APO) at 64,000× magnification.** The flowchart illustrates the image-processing workflow initiated from 3,301 movie stacks using the super-resolution (Bin1) strategy. Following patch motion correction and CTF estimation, 3,301 micrographs were obtained, yielding 4,905,391 template-picked particles. After 2D classification, 2,117,987 particles were retained for *ab-initio* reconstruction and heterogeneous refinement. The predominant 3D class (outlined in red) accounted for ~92.6% of the particles, yielding 1,957,152 particles after extraction (Box size: 420, Bin1) that were parallelly used for Bin1 and downsampled Bin2 (Box size: 210) refinements. Prior to reference-based motion correction, homogeneous refinement yielded resolutions of 1.86 Å (Bin1) and 2.74 Å (Bin2), respectively. Following reference-based motion correction, the final reconstructions achieved 1.72 Å for Bin1 (1,957,082 particles) and 2.74 Å for Bin2 (1,957,073 particles). Each local-resolution map (contour at 4 σ) features a color scale on the right and the global resolution noted below.

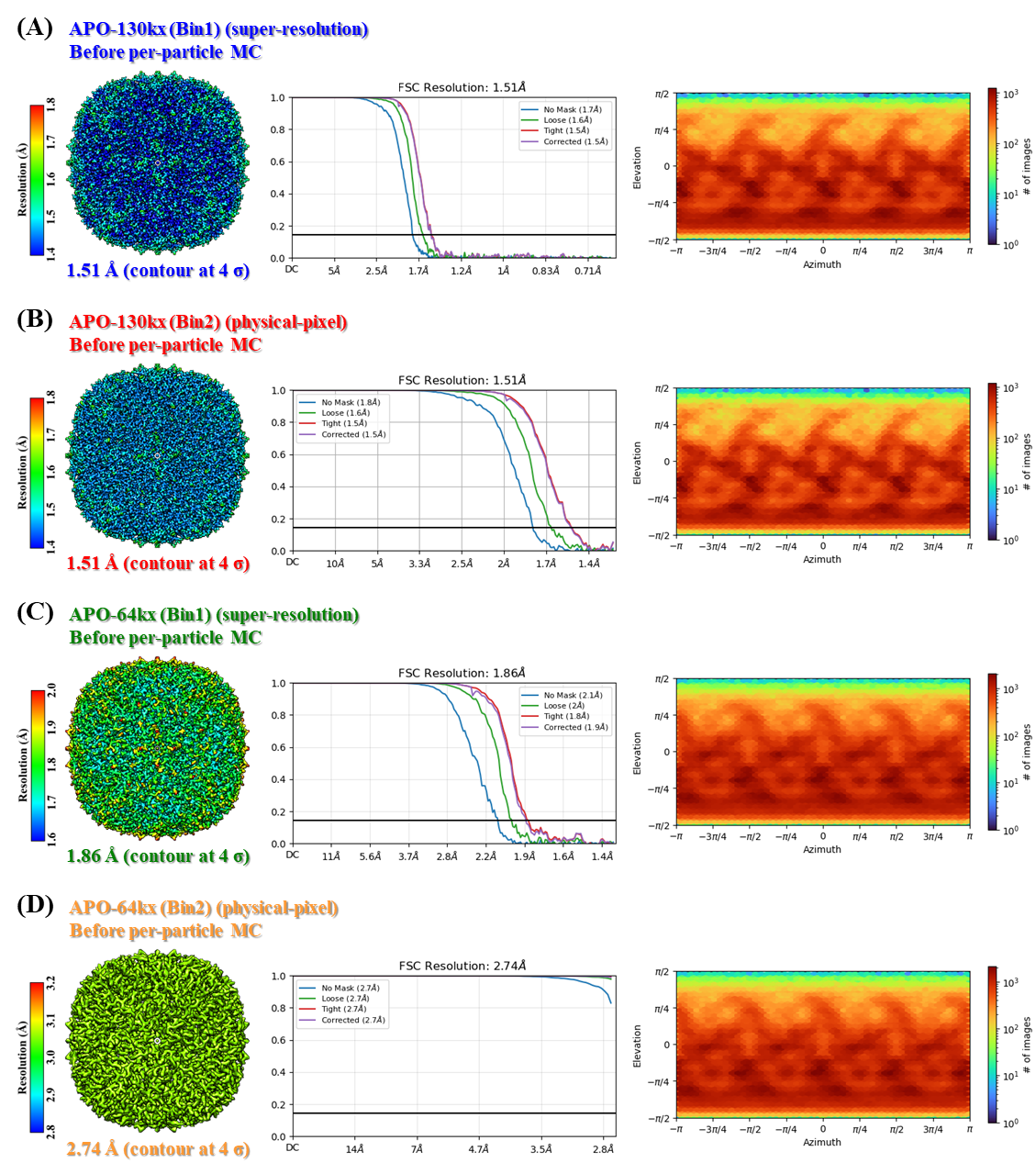

**Figure S4. Reconstruction quality assessment of apoferritin (APO) before per-particle motion correction.** Local-resolution maps (left), gold-standard FSC curves (middle), and particle-orientation distribution plots (right) are systematically compared for reconstructions under different imaging and sampling conditions: **(A)** 130,000× Bin1 (super-resolution, 1.51 Å), **(B)** 130,000× Bin2 (physical-pixel, 1.51 Å), **(C)** 64,000× Bin1 (super-resolution, 1.86 Å), and **(D)** 64,000× Bin2 (physical-pixel, 2.74 Å). Each local-resolution map, displayed at a contour level of 4 σ, features a color scale on its left, with the corresponding FSC curve and global resolution indicated in the center. Orientation plots on the right illustrate particle angular coverage, with their respective color scales representing particle counts across orientations.

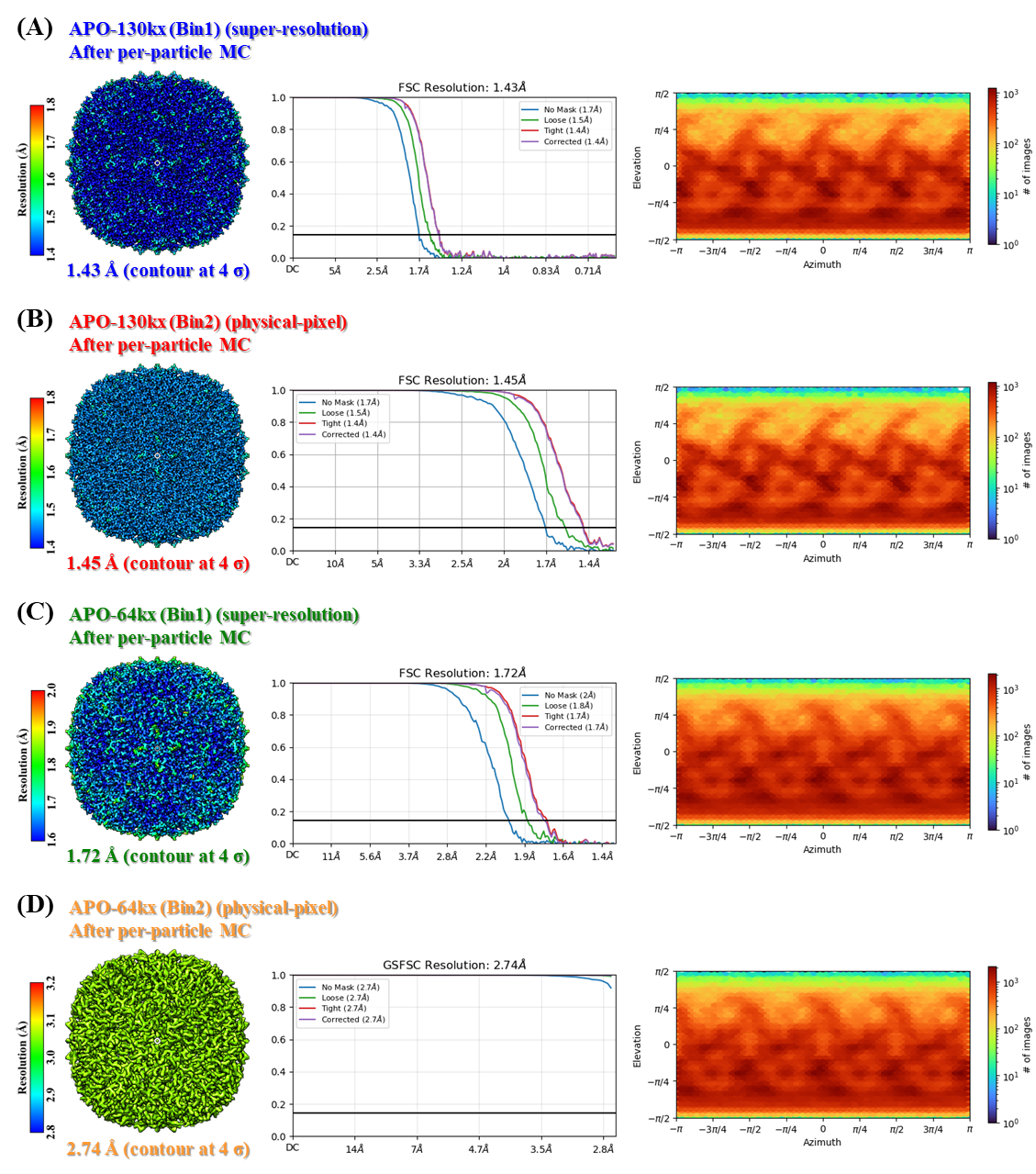

**Figure S5. Reconstruction quality assessment of apoferritin (APO) after per-particle motion correction.** Local-resolution maps (left), gold-standard FSC curves (middle), and particle-orientation distribution plots (right) are systematically compared for reconstructions under different imaging and sampling conditions: **(A)** 130,000× Bin1 (super-resolution, 1.43 Å), **(B)** 130,000× Bin2 (physical-pixel, 1.45 Å), **(C)** 64,000× Bin1 (super-resolution, 1.72 Å), and **(D)** 64,000× Bin2 (physical-pixel, 2.74 Å). Each local-resolution map, displayed at a contour level of 4 σ, features a color scale on its left, with the corresponding FSC curve and global resolution indicated in the center. Orientation plots on the right illustrate particle angular coverage, with their respective color scales representing particle counts across orientations.

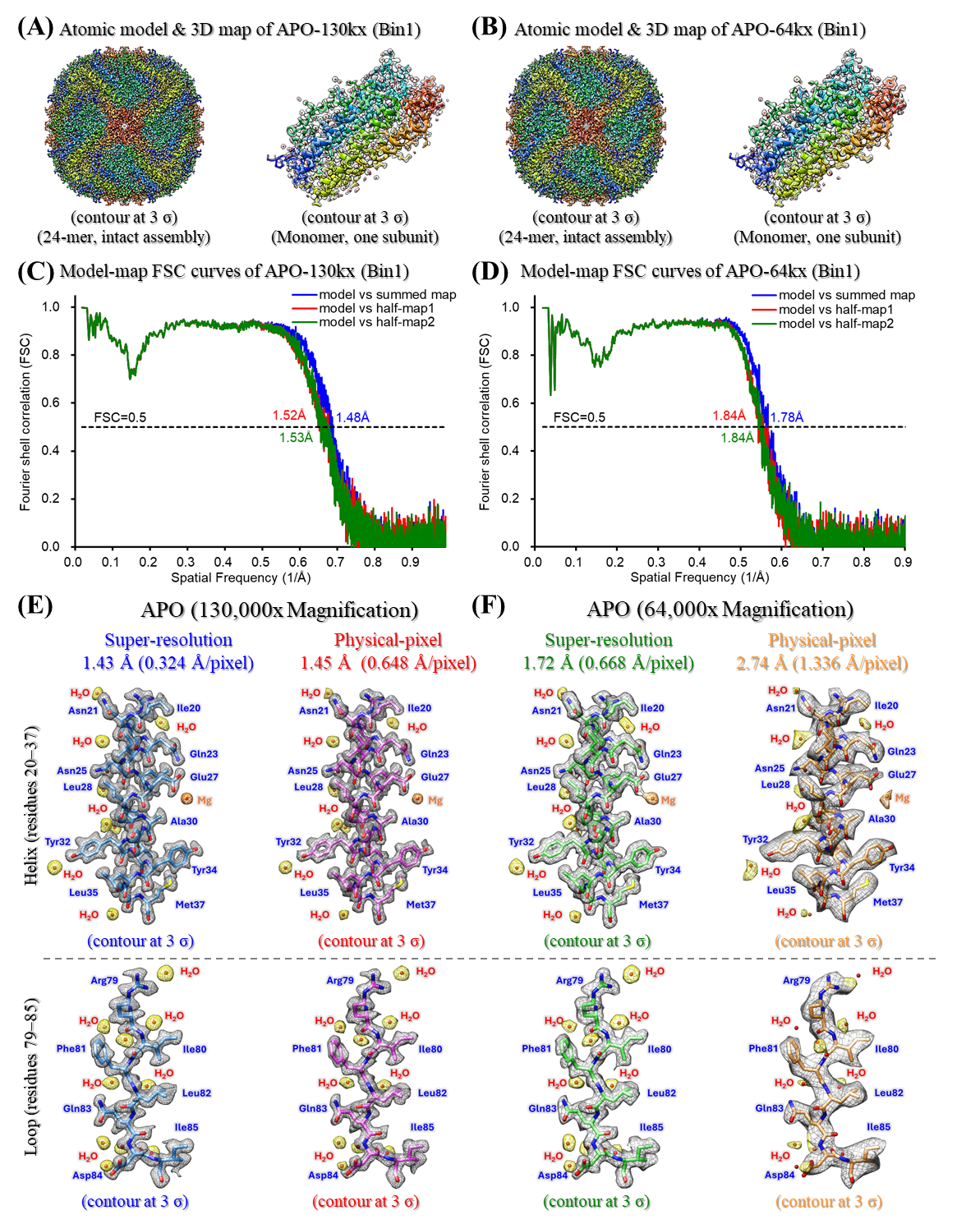

**Figure S6. Validation of apoferritin (APO) atomic models and local detail preservation across magnifications and sampling strategies. (A–B)** Three-dimensional density maps (transparent gray, contoured at 3 σ) are overlaid with refined atomic models of APO reconstructed at nominal magnifications of **(A)** 130,000× and **(B)** 64,000×. Both the intact 24-subunit assembly with octahedral symmetry and a representative subunit are displayed, with residues rainbow-colored sequentially from the N-terminus (blue) to the C-terminus (red). **(C–D)** Model-to-map Fourier shell correlation (FSC) curves for the **(C)** 130,000× and **(D)** 64,000× datasets. Cross-validation curves represent the atomic model versus the full summed map (blue), work half-map 1 (red), and free half-map 2 (green), with the standard FSC = 0.5 validation threshold indicated by the horizontal dashed line. **(E–F)** Parallel comparisons of local structural detail preservation within the alpha-helix (residues 20–37, top row, contoured at 3 σ) and loop (residues 79–85, bottom row, contoured at 3 σ) regions under **(E)** 130,000× and **(F)** 64,000× workflows. Global map resolutions and pixel sizes for each condition are annotated at the top of each column. Density maps are rendered as transparent gray meshes overlaid with the corresponding atomic models shown as colored sticks. These contour thresholds were selected to optimally visualize structural features for each reconstruction. Coordinated water molecules (H₂O) and magnesium ions (Mg²⁺) are represented as red and orange spheres, respectively.

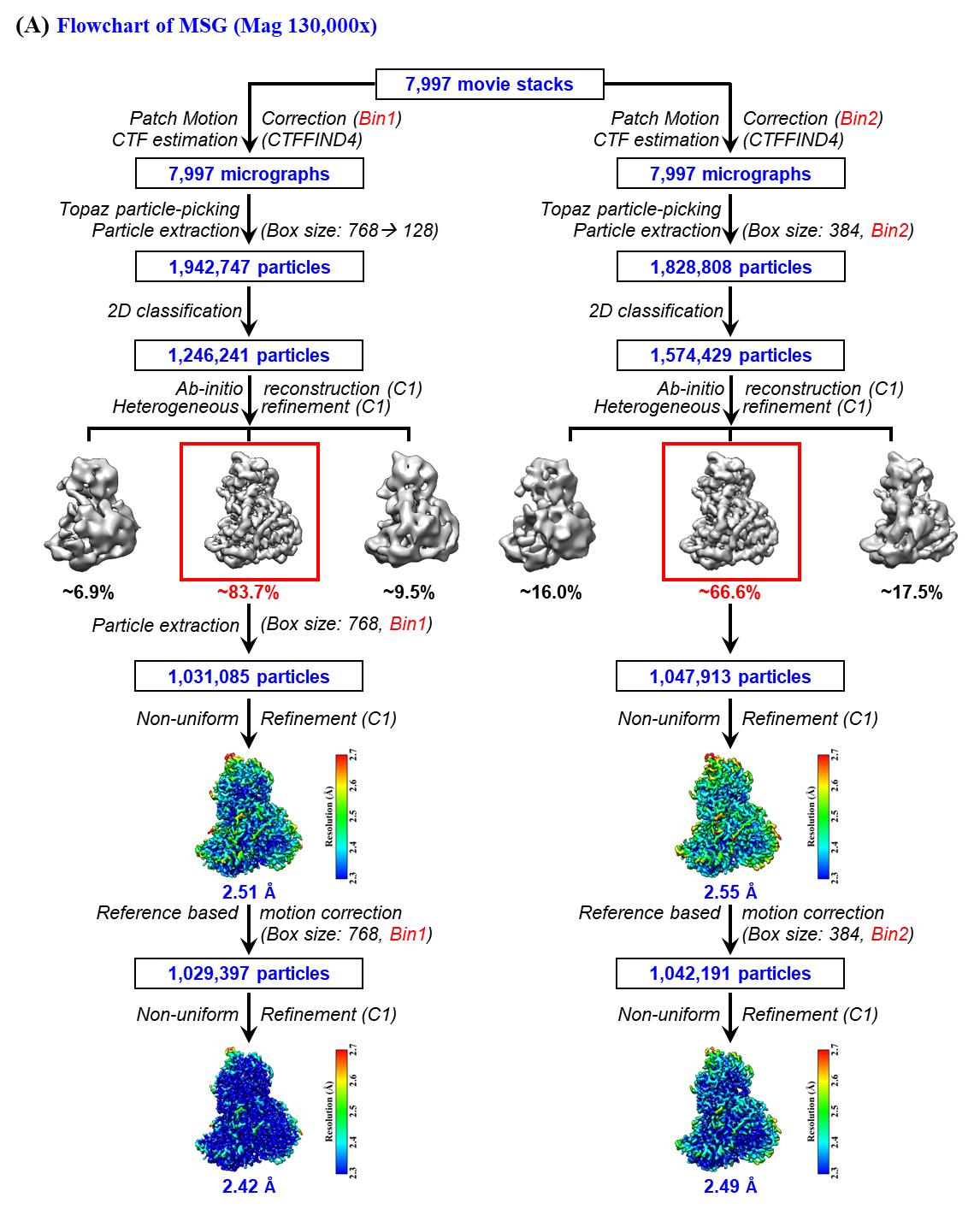

**Figure S7. Cryo-EM data processing workflow for malate synthase G (MSG) at 130,000× magnification.** The flowchart illustrates the parallel processing strategies derived from the same 7,997 movie stacks using super-resolution (Bin1, left) and physical-pixel (Bin2, right) workflows. Topaz-based particle picking initially yielded 1,942,747 (Bin1) and 1,828,808 (Bin2) particles. Following 2D classification, 1,246,241 (Bin1) and 1,574,429 (Bin2) particles were selected for *ab-initio* reconstruction and heterogeneous refinement in C1 symmetry. The predominant 3D class (outlined in red) accounted for ~83.7% (Bin1) and ~66.6% (Bin2) of the particles, resulting in 1,031,085 and 1,047,913 particles after extraction, respectively. Prior to reference-based motion correction, non-uniform refinement yielded global resolutions of 2.51 Å (Bin1) and 2.55 Å (Bin2). Following reference-based motion correction, the final reconstructions using 1,029,397 (Bin1) and 1,042,191 (Bin2) particles achieved resolutions of 2.42 Å and 2.49 Å, respectively. Each local-resolution map (contour at 8 σ) features a color scale on the right and the global resolution noted below.

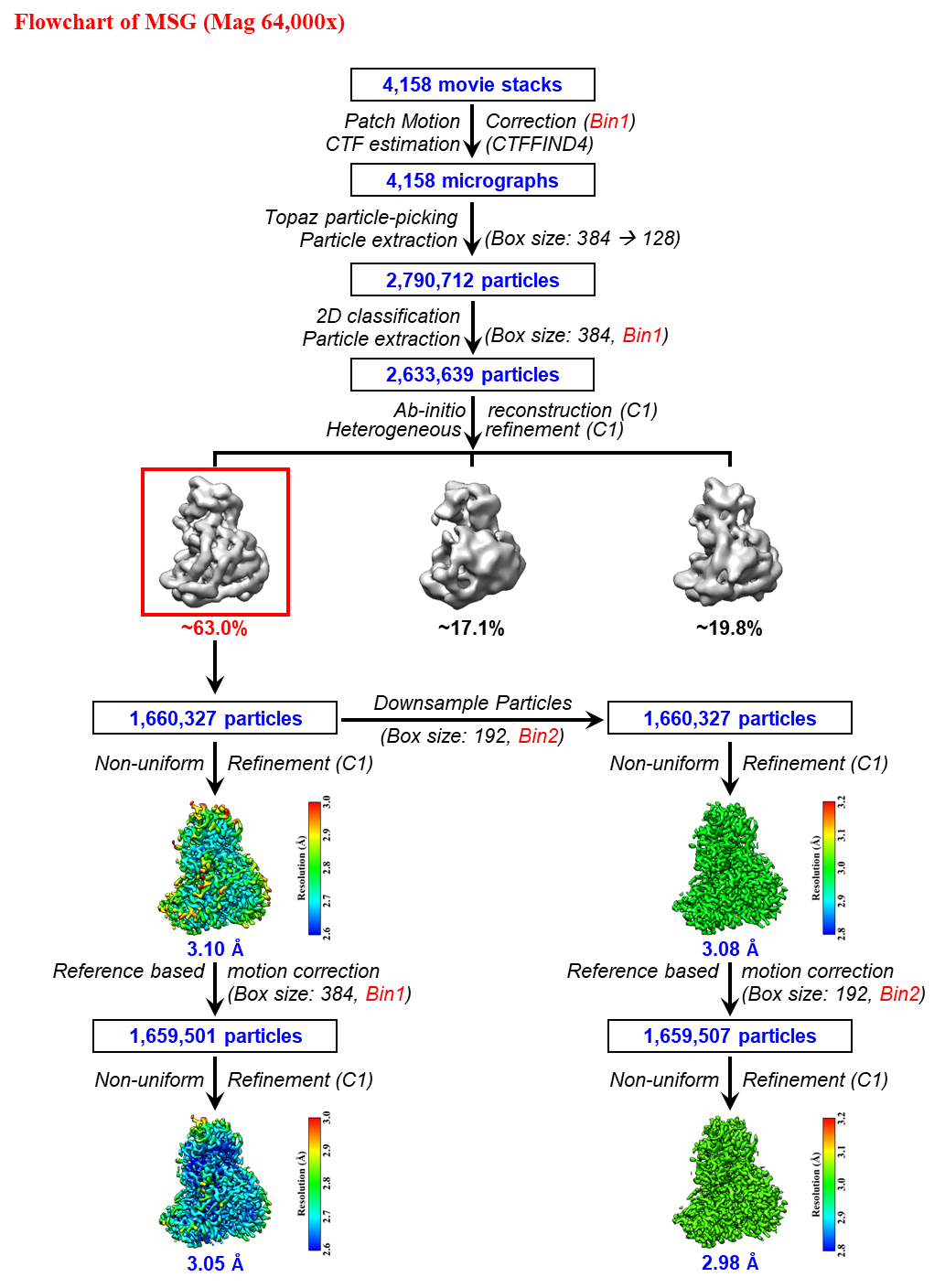

**Figure S8. Cryo-EM data processing workflow for malate synthase G (MSG) at 64,000× magnification.** The flowchart illustrates the image-processing workflow initiated from 4,158 movie stacks using the super-resolution (Bin1) strategy. Following patch motion correction and CTF estimation, 4,158 micrographs were obtained, from which 2,790,712 particles were initially picked using Topaz. After 2D classification, 2,633,639 particles were retained for *ab-initio* reconstruction and heterogeneous refinement in C1 symmetry. The predominant 3D class (outlined in red) accounted for ~63.0% of the particles, yielding 1,660,327 particles after extraction (Box size: 384, Bin1) that were used for non-uniform refinement and subsequently downsampled for parallel Bin2 (Box size: 192) analysis. Prior to reference-based motion correction, non-uniform refinement yielded global resolutions of 3.10 Å (Bin1) and 3.08 Å (Bin2). Following reference-based motion correction, the final reconstructions achieved 3.05 Å for Bin1 (1,659,501 particles) and 2.98 Å for Bin2 (1,659,507 particles). Each local-resolution map (contour at 8 σ) features a color scale on the right and the global resolution noted below.

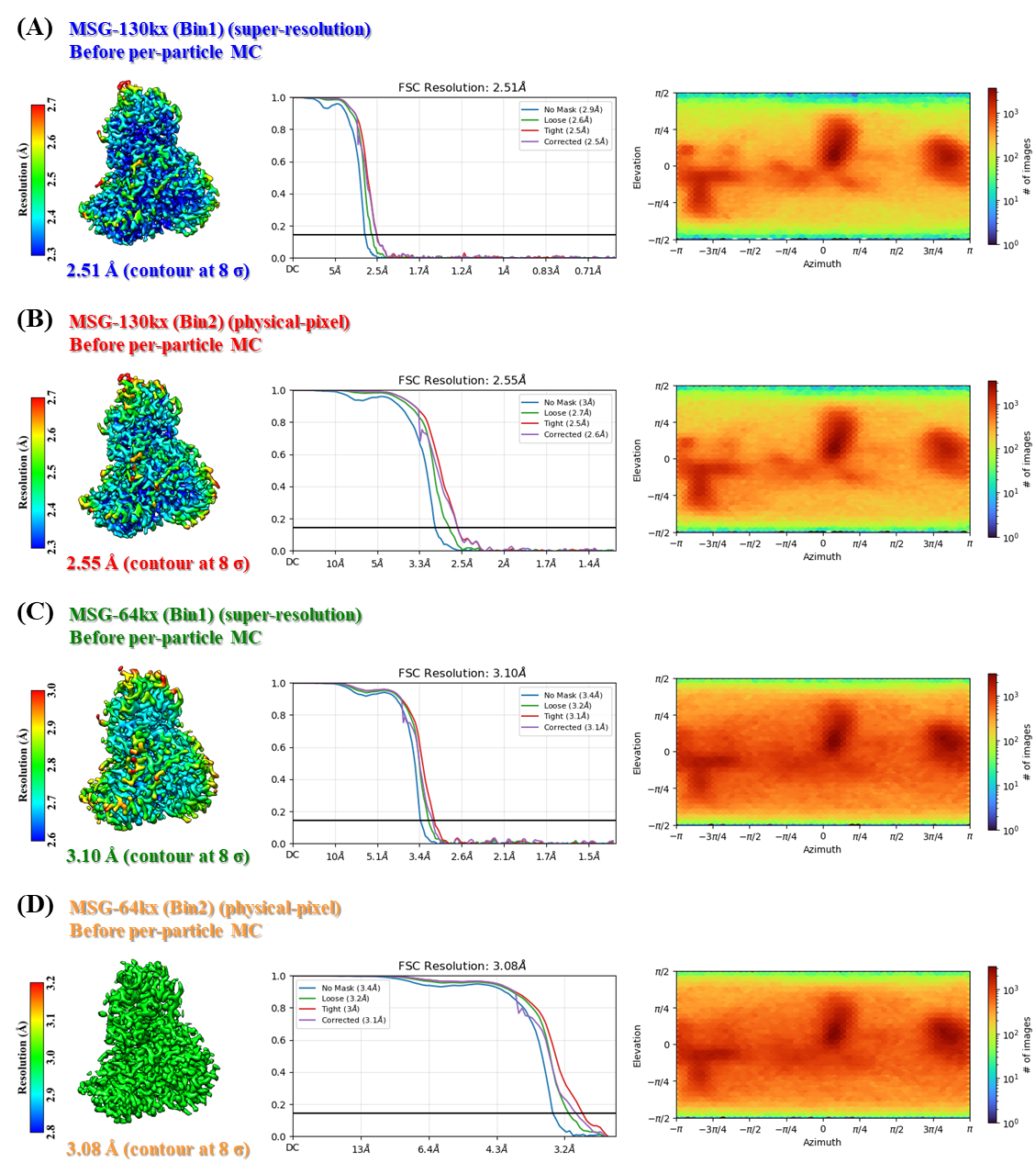

**Figure S9. Reconstruction quality assessment of malate synthase G (MSG) before per-particle motion correction.** Local-resolution maps (left), gold-standard FSC curves (middle), and particle-orientation distribution plots (right) are systematically compared for reconstructions under different imaging and sampling conditions: **(A)** 130,000× Bin1 (super-resolution, 2.51 Å), **(B)** 130,000× Bin2 (physical-pixel, 2.55 Å), **(C)** 64,000× Bin1 (super-resolution, 3.10 Å), and **(D)** 64,000× Bin2 (physical-pixel, 3.08 Å). Each local-resolution map, displayed at a contour level of 8 σ, features a color scale on its left, with the corresponding FSC curve and global resolution indicated in the center. Orientation plots on the right illustrate particle angular coverage, with their respective color scales representing particle counts across orientations.

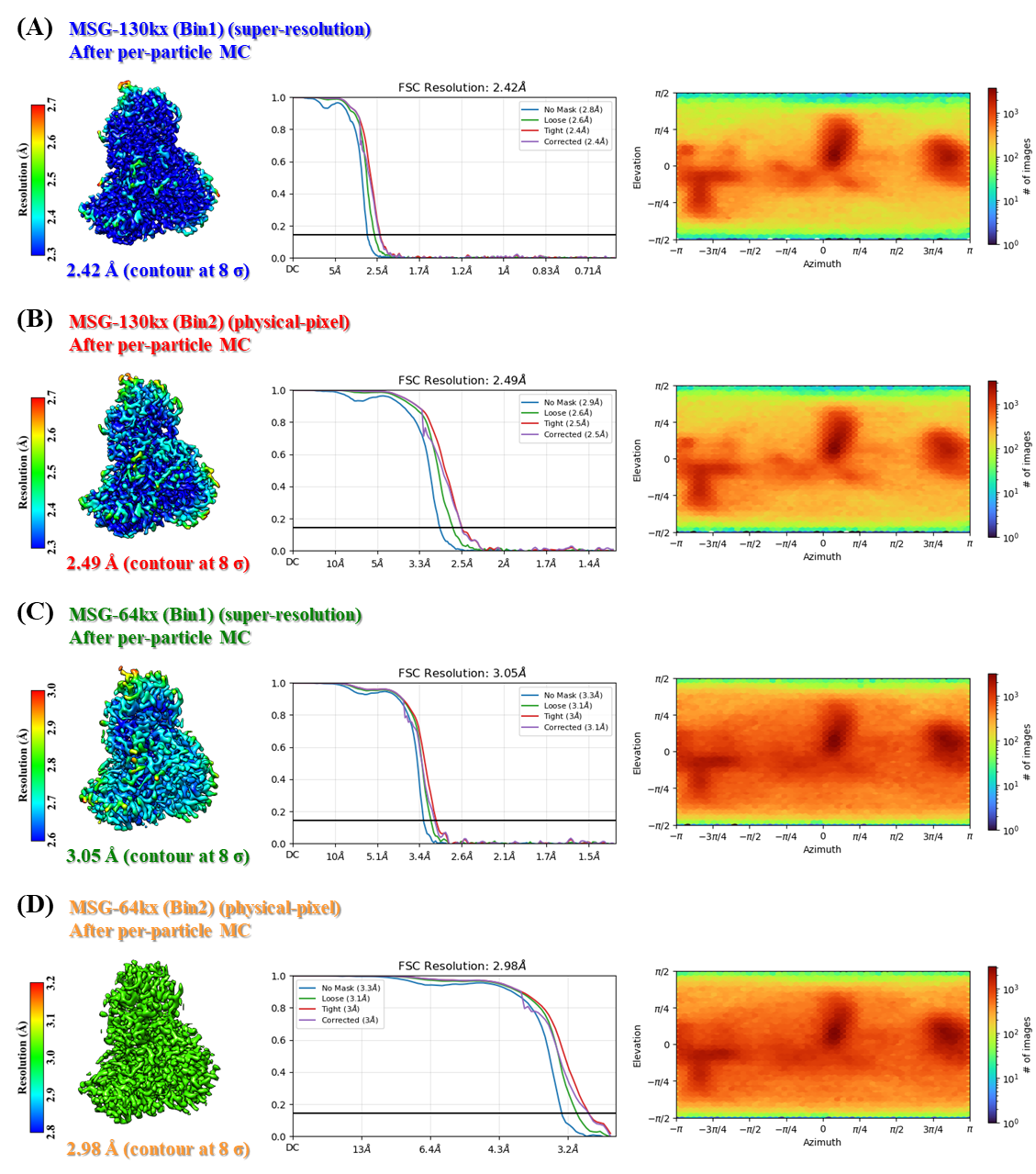

**Figure S10. Reconstruction quality assessment of malate synthase G (MSG) after per-particle motion correction.** Local-resolution maps (left), gold-standard FSC curves (middle), and particle-orientation distribution plots (right) are systematically compared for reconstructions under different imaging and sampling conditions: **(A)** 130,000× Bin1 (super-resolution, 2.42 Å), **(B)** 130,000× Bin2 (physical-pixel, 2.49 Å), **(C)** 64,000× Bin1 (super-resolution, 3.05 Å), and **(D)** 64,000× Bin2 (physical-pixel, 2.98 Å). Each local-resolution map, displayed at a contour level of 8 σ, features a color scale on its left, with the corresponding FSC curve and global resolution indicated in the center. Orientation plots on the right illustrate particle angular coverage, with their respective color scales representing particle counts across orientations.

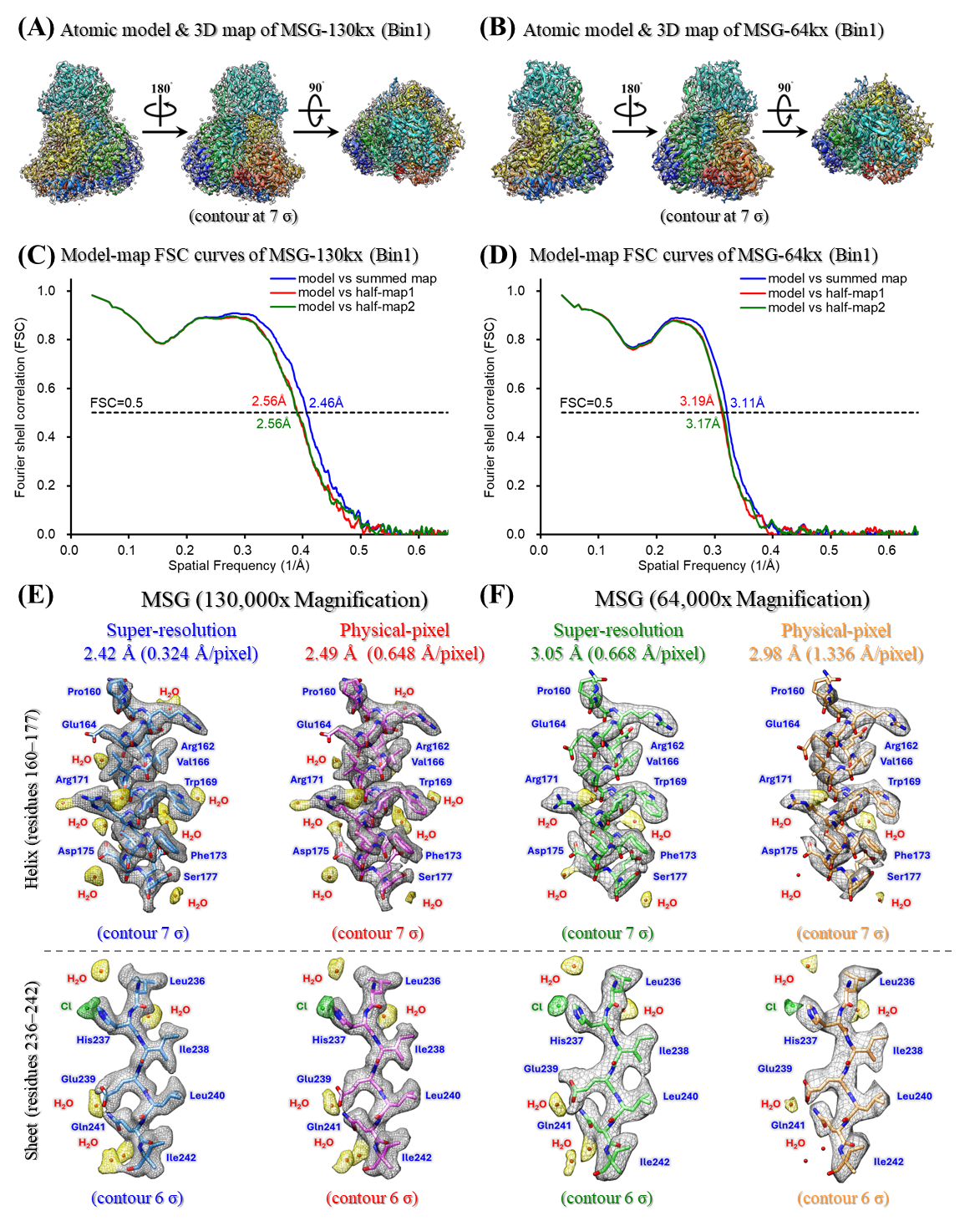

**Figure S11. Validation of malate synthase G (MSG) atomic models and local detail preservation across magnifications and sampling strategies. (A–B)** Three-dimensional density maps (transparent gray, contoured at 7 σ) are overlaid with refined atomic models of MSG reconstructed at nominal magnifications of **(A)** 130,000× and **(B)** 64,000×. Multiple orientations, including 90° and 180° rotations, are displayed to evaluate the global monomeric fit, with residues rainbow-colored sequentially from the N-terminus (blue) to the C-terminus (red). **(C–D)** Model-to-map Fourier shell correlation (FSC) curves for the **(C)** 130,000× and **(D)** 64,000× datasets. Cross-validation curves represent the atomic model versus the full summed map (blue), work half-map 1 (red), and free half-map 2 (green), with the standard FSC = 0.5 validation threshold indicated by the horizontal dashed line. **(E–F)** Parallel comparisons of local structural detail preservation within the alpha-helix (residues 160–177, top row, contoured at 7 σ) and beta-sheet (residues 236–242, bottom row, contoured at 6 σ) regions under **(E)** 130,000× and **(F)** 64,000× workflows. Global map resolutions and pixel sizes for each condition are annotated at the top of each column. Density maps are rendered as transparent gray meshes overlaid with the corresponding atomic models shown as colored sticks. These contour thresholds were selected to optimally visualize structural features for each reconstruction. Coordinated water molecules (H₂O) and chloride ions (Cl⁻) are represented as red and green spheres, respectively.

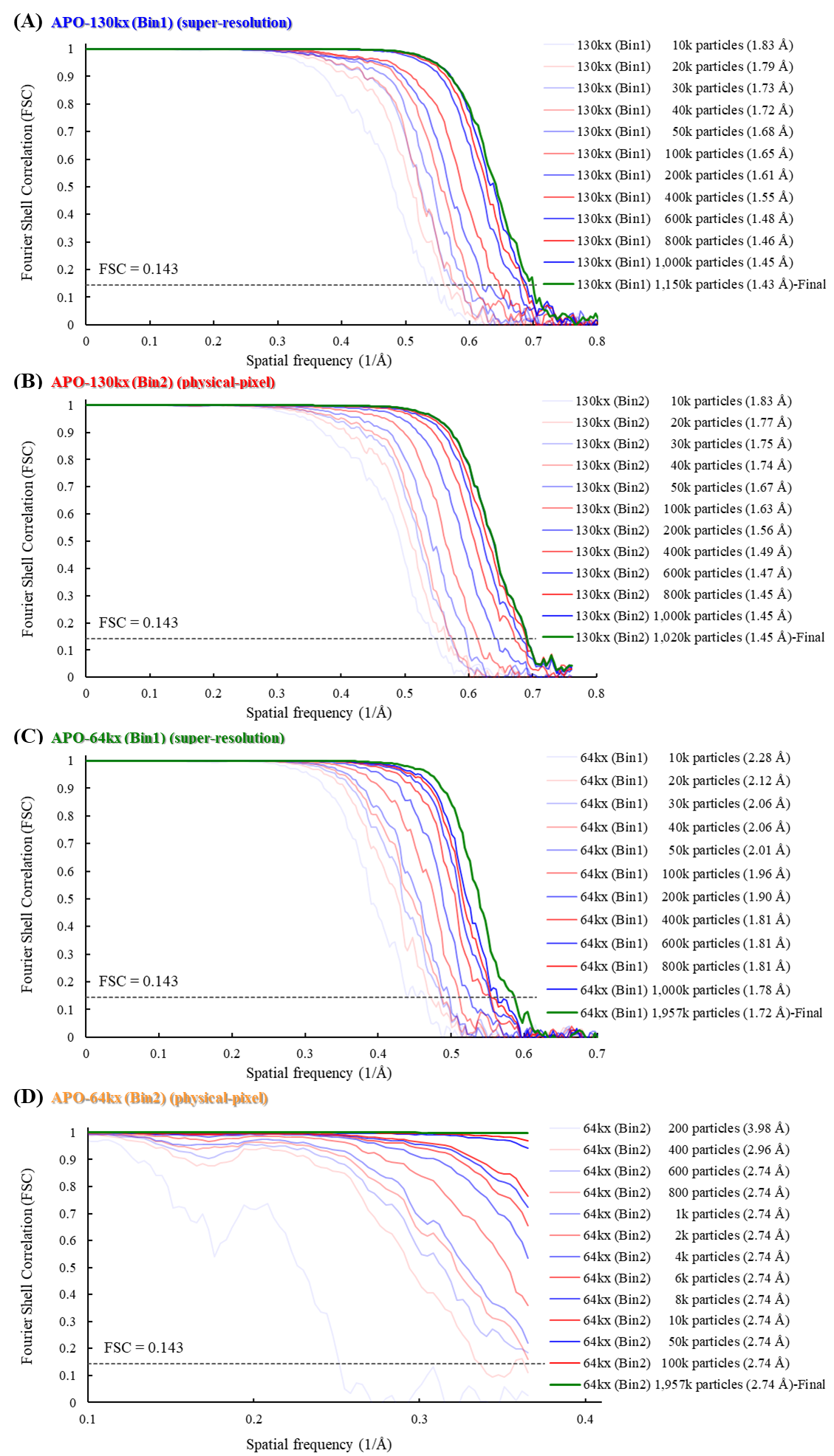

**Figure S12. Gold-standard FSC curves for apoferritin (APO) reconstructions across varying particle numbers.** Gold-standard Fourier shell correlation (FSC) curves are systematically compared for reconstructions under different imaging and sampling conditions: **(A)** 130,000× Bin1 (super-resolution), with particle subsets ranging from 10,000 to the final 1,150,038 particles, yielding resolutions from 1.83 Å to 1.43 Å; **(B)** 130,000× Bin2 (physical-pixel), with subsets from 10,000 to the final 1,020,042 particles, yielding resolutions from 1.83 Å to 1.45 Å; **(C)** 64,000× Bin1 (super-resolution), with subsets from 10,000 to the final 1,957,082 particles, yielding resolutions from 2.28 Å to 1.72 Å; and **(D)** 64,000× Bin2 (physical-pixel), with subsets from 200 to the final 1,957,073 particles, yielding resolutions from 3.98 Å to 2.74 Å. Global map resolutions were determined using the gold-standard FSC = 0.143 criterion (horizontal dashed line). All reconstructions were performed following reference-based motion correction. The color gradient from light to dark denotes an increasing number of particles, with the final reconstruction curves highlighted in bold green.

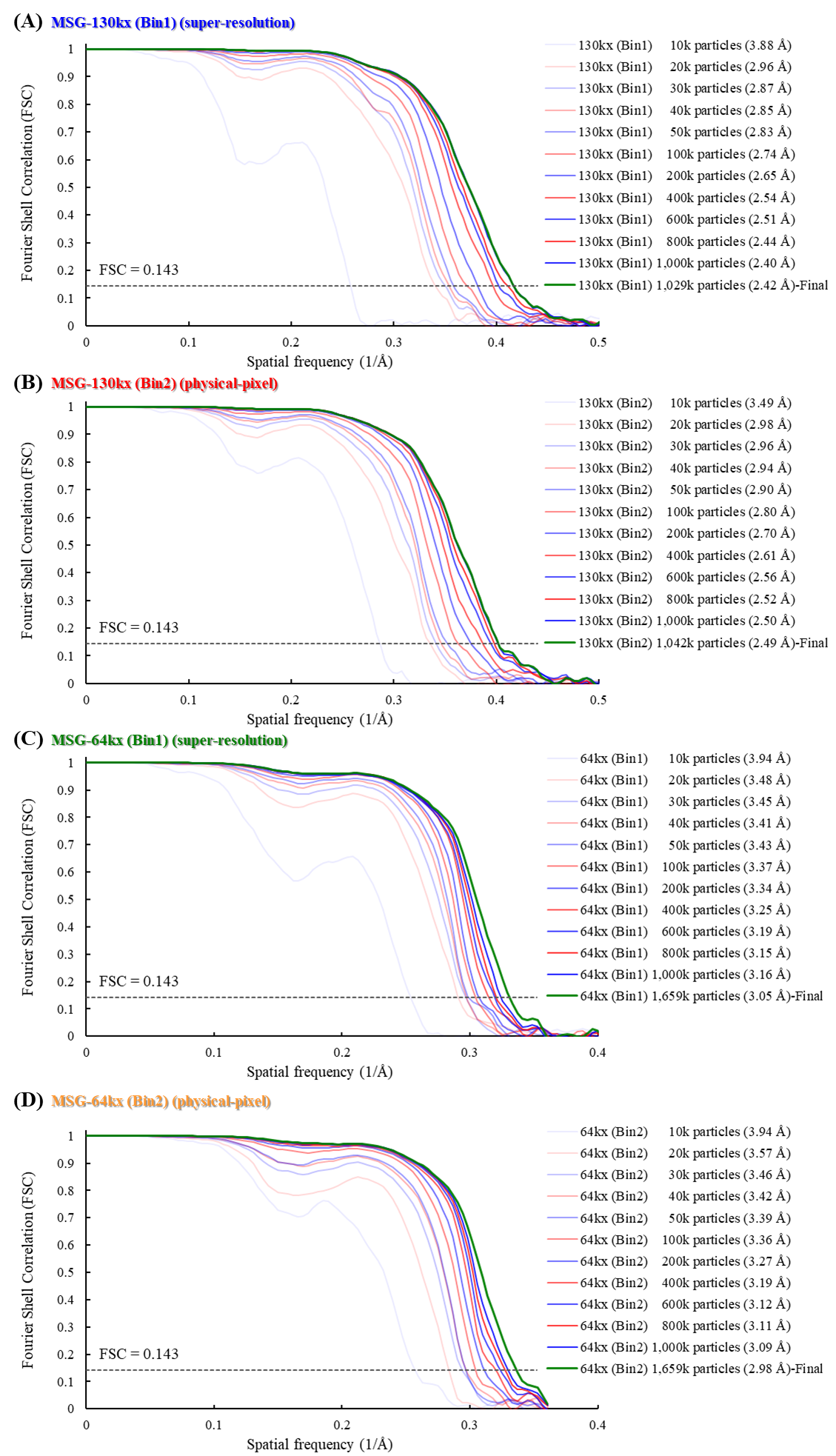

**Figure S13. Gold-standard FSC curves for malate synthase G (MSG) reconstructions across varying particle numbers.** Gold-standard Fourier shell correlation (FSC) curves are systematically compared for reconstructions under different imaging and sampling conditions: **(A)** 130,000× Bin1 (super-resolution), with particle subsets ranging from 10,000 to the final 1,029,397 particles, yielding resolutions from 3.88 Å to 2.42 Å; **(B)** 130,000× Bin2 (physical-pixel), with subsets from 10,000 to the final 1,042,191 particles, yielding resolutions from 3.49 Å to 2.49 Å; **(C)** 64,000× Bin1 (super-resolution), with subsets from 10,000 to the final 1,659,501 particles, yielding resolutions from 3.94 Å to 3.05 Å; and **(D)** 64,000× Bin2 (physical-pixel), with subsets from 10,000 to the final 1,659,507 particles, yielding resolutions from 3.94 Å to 2.98 Å. Global map resolutions were determined using the gold-standard FSC = 0.143 criterion (horizontal dashed line). All reconstructions were performed following reference-based motion correction. The color gradient from light to dark denotes an increasing number of particles, with the final reconstruction curves highlighted in bold green.
